# Sparse connectivity for MAP inference in linear models using sister mitral cells

**DOI:** 10.1101/2021.06.28.450144

**Authors:** Sina Tootoonian, Andreas Schaefer, Peter Latham

**Affiliations:** Gatsby Computational Neuroscience Unit, University College London, London, UK; Sensory Circuits and Neurotechnology Laboratory, The Francis Crick Institute, London, UK

## Abstract

Sensory processing is hard because the variables of interest are encoded in spike trains in a relatively complex way. A major goal in studies of sensory processing is to understand how the brain extracts those variables. Here we revisit a common encoding model in which variables are encoded linearly. Although there are typically more variables than neurons, this problem is still solvable because only a small number of variables appear at any one time (sparse prior). However, previous solutions require all-to-all connectivity, inconsistent with the sparse connectivity seen in the brain. Here we propose an algorithm that provably reaches the MAP (maximum *a posteriori*) inference solution, but does so using sparse connectivity. Our algorithm is inspired by the circuit of the mouse olfactory bulb, but our approach is general enough to apply to other modalities. In addition, it should be possible to extend it to nonlinear encoding models.

**Summary:** Sensory systems must infer latent variables from noisy and ambiguous input. MAP inference – choosing the most likely latent variable given the sensory input – is one of the simplest methods for doing that, but its neural implementation often requires all-to-all connectivity between the neurons involved. In common sensory contexts this can require a single neuron to connect to hundreds of thousands of others, which is biologically implausible. In this work we take inspiration from the ‘sister’ mitral cells of the olfactory system – groups of neurons associated with the same input channel – to derive a method for performing MAP inference using sparse connectivity. We do so by assigning sister cells to random subsets of the latent variables and using additional cells to ensure that sisters correctly share information. We then derive the circuitry and dynamics required for the sister cells to compute the original MAP inference solution. Our work yields a biologically plausible circuit that provably solves the MAP inference problem and provides experimentally testable predictions. While inspired by the olfactory system, our method is quite general, and is likely to apply to other sensory modalities.

## 1 Introduction

A common view of sensory systems is that they invert generative models of the environment to infer the causes underlying sensory input. Sensory input is typically ambiguous, so a given input can be explained by multiple causes. Consequently, correct inference requires adequately accounting for interactions among causes. For example, increased evidence for one cause often reduces the probability of, or “explains away”, competing causes (if you think the object you’re smelling is an orange, that makes it less likely to be a lemon). Any neural circuit performing inference must therefore implement mechanisms for inter-causal interaction. This typically results in dense – and in many case all-to-all – connectivity between neurons representing causes. The myriad causes potentially responsible for a given sensory input often require a neuron representing a cause to connect to hundreds of thousands of others. Such dense connectivity is biologically implausible.

This problem is easy to demonstrate in linear models of sensory input. (Although linear may seem overly restrictive, in fact such models have been successful in explaining basic features of the visual [1], olfactory [2], and auditory [3] systems). Consider noisy receptors *y_i_* (e.g. retinal ganglion cells, olfactory glomeruli) linearly excited by causes *x_j_* (e.g. edges, odours) according to a matrix *A_ij_*. Under this model, the excitation of the *i*^th^ receptor is given by

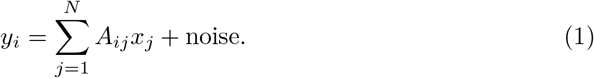

The causes responsible for the observations, *y_i_*, can be estimated by minimizing, for example, the squared error between the *actual* observations and the *expected* observations, subject to a penalty on the magnitude of the inferred causes *x_j_*. A population of neurons whose individual firing rates represent the *x_j_* can do this by gradient descent [4],

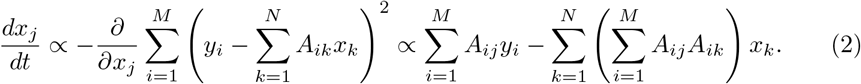

These dynamics can be interpreted as balancing the evidence for the cause *x_j_* due to the receptor inputs *y_i_* (the first term) while accounting for the explanatory power of the other causes (the second term). In particular, 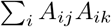 reflects the contribution of cause *x_k_* to the evidence for cause *x_j_*. Importantly, even if the elements *A_ij_* are sparse, 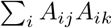 will be non-zero for most *j* and *k*, implying nearly all-all connectivity in a circuit implementing Eq. (2). In common sensory settings there may be hundreds of thousands of causes that explain a given input. This means that *x_j_* must connect to hundreds of thousands of other neurons.

Below we show how the problem of all-to-all connectivity can be solved so that inference can be performed with realistically sparse connectivity. We begin by recapitulating the MAP inference problem, focusing on the olfactory setting for concreteness. This is basically sparse coding applied to olfaction, and suffers from all-to-all connectivity. We then derive a solution inspired by the anatomy of the vertebrate olfactory bulb, namely the presence of dozens of ‘sister cells’ that receive input from the same glomerulus. That solution leads to MAP inference, but using sparser connectivity. While we focus here on the olfactory system, our method is applicable to other modalities.

## 2 Results

### 2.1 Olfaction as MAP inference

Animals observe odours indirectly via the excitation of olfactory receptor neurons that project their axons to spherical bundles of neuropil called glomeruli. Each receptor neuron is thought to express a single receptor gene [5] from a large repertoire, and neurons expressing the same gene almost always converge onto one of two glomeruli, on either side of the olfactory bulb [6]. Thus each glomerulus represents the pooled activity of the receptor neurons expressing a single type of olfactory receptor. We represent this vector of glomerular activations by **y** = {*y*_1_, *y*_2_,…, *y_M_*}, where *y_i_* is the activation of the *i*’th glomerulus, and *M* is the number of glomeruli per lobe of the olfactory bulb, or equivalently, the number of olfactory receptor genes expressed by the animal. This number is ~50 for flies [7], ~300 for humans [8], and ~1000 for mice [9].

The task of the animal is to infer the odour, **x** (which consists of *N* components, {*x*_1_, *x*_2_,…, *x_N_*}), from the receptor activations, **y** (see Fig. 1A). There are two main interpretations for the *x_j_*. One is that *x_j_* is the concentration of the *j*^th^ molecule in the odour, and so *N* is the number of distinct molecular species that the animal may encounter in its environment. The other is that *x_j_* represents a complete olfactory object (e.g., coffee, bacon, marmalade) rather than a molecular species; in this case, *N* is the number of learned odours. To estimate *N* for the first interpretation, we note that the study of an estimated 0.25% of all flowering plants has yielded 1700 floral scent compounds [10], suggesting an upper estimate for *N* on the order of 1700/0.0025, or roughly 700,000 (though the actual number could turn out to be far fewer if existing molecules appear in as-yet-undiscovered floral scents), on the same order as the 400,000 estimated in the literature [11]. For the second interpretation (odours are complex olfactory objects), *N* is difficult to approximate, but estimates for the number of *distinguishable* odour objects range from 10,000 [12] to 1 trillion [13]. Here we simply assume that in both cases *N* is large. In either case, odours are very sparse – either because only a few molecular species are present [14], or only a few odours are present.

**Fig 1.**
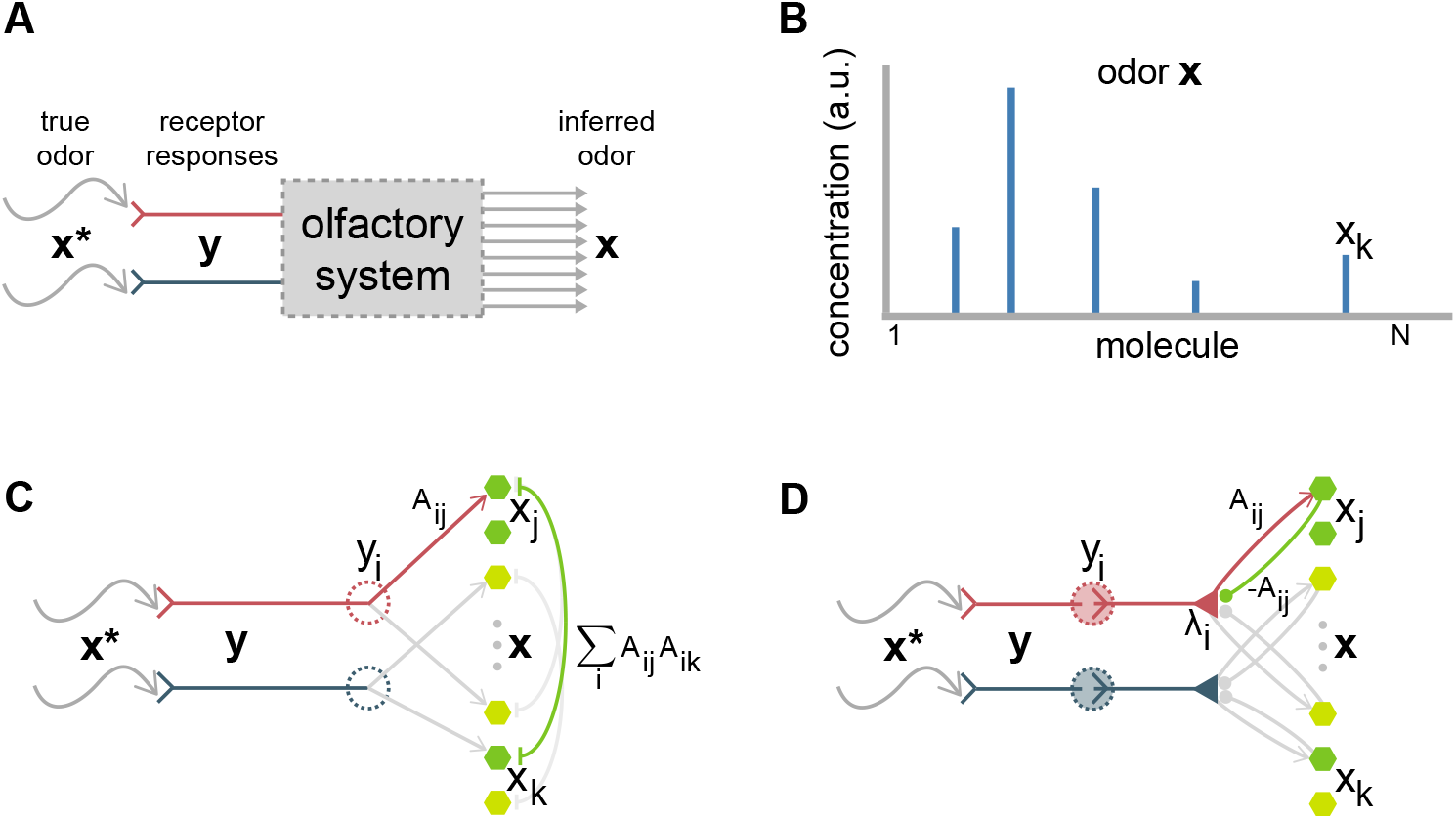
Olfaction as MAP inference. (A) Animals observe odours **x**\* indirectly via receptor activations. We assume the function of the olfactory system is to report the odour **x** most likely to have caused the observed receptor activations **y**. (B) Schematic representation of an odour, whose defining feature is that it’s sparse (meaning very few components are active). (C) A basic circuit for performing MAP inference on odours: Receptor *i* projects directly to each readout unit *x_j_* with weight *A_ij_* determined by the affinity of receptor *i* for molecule *j*; the readout unit *x_j_* reciprocally inhibits unit *x_k_* with weight 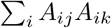. The latter term is likely be non-zero even if *A_ij_* is sparse (since that requires only one term in the sum over *i* to be nonzero), resulting in each readout unit inhibiting and being inhibited by potentially ~100,000 other units. (D) An alternate circuit that performs the same inference. Mitral cells now mediate between glomeruli and the readout units. Each mitral cell *λ_i_* excites each readout unit *x_j_* with weight *A_ij_* and is in turn inhibited by the same amount. No inhibition is needed between readout units, but a mitral cell must still excite and be inhibited by each of potentially ~100,000 readout units.

From the point of view of learning and inference, there are advantages to both interpretations. If odours are represented as molecular concentration vectors, then information about how each molecule excites each receptor is determined only by the physical parameters of the molecule and receptor. It can, therefore, be learned on evolutionary timescales and hard-wired into the circuit, at least in principle, and it provides a simple substrate for the animal to generalize between chemically similar odours. It is disadvantageous in that the animal requires higher-order circuitry to infer learned olfactory objects (e.g., coffee, bacon, marmalade), which consist of many types of odour molecules. If odours are represented as complex objects, then those objects have to be learned within the lifetime of the animal. However, once learned, further higher order circuitry is not needed. Although these important representational issues are beyond the scope of the work presented here, we mention them in passing as examples of the non-trivial assumptions required before a theory of olfactory circuit function can be developed.

We assume a very simple model of the transduction of odours into neural activations: odour components contribute linearly to the input current of a receptor, which is then converted into a firing rate by a static, invertible, point-wise non-linearity. That is, the excitation of glomerulus *i* is described as

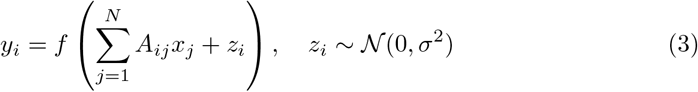

where *x_j_* is the concentration of the *j*^th^ molecule, *A_ij_* is the *affinity* of the *i*^th^ receptor for the *j*’th molecule, *f* is a static pointwise invertible nonlinearity converting input current to firing rate, and *z_i_* is additive noise with variance *σ*^2^. Such nonlinearities can be inverted without changing the nature of the inference problem, so, without loss of generality, we take *f* to be the identity. Thus, our likelihood is

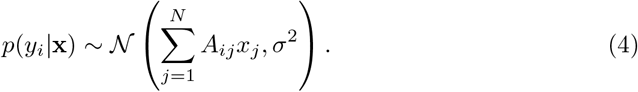

Because the number of glomeruli, *M*, is likely to be much smaller than the number of molecular species, *N*, a whole manifold of odours can be consistent with any particular pattern of glomerular activation. We resolve this ambiguity by selecting the candidate odour that is most consistent with our prior information about odours. The prior *p*(**x**) encodes the animal’s background knowledge about the presence of odours in the environment. We assume that the animal makes the simplifying assumptions that molecules appear independently of each other (but see [15], [16]), and that the marginal probability distribution for each molecule, *p*(*x_i_*), has the same form.

We use an elastic net distribution (a combination of *ℓ*_1_ and *ℓ*_2_ penalties) [17,18]. The *ℓ*_1_ penalty promotes sparsity as is observed [14] (see Fig. 1B); the *ℓ*_2_ penalty discourages very large concentrations. In addition, we include a term enforcing the non-negativity of concentrations, yielding

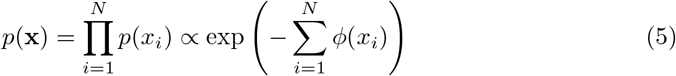

where

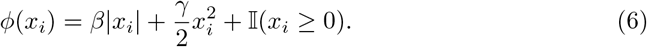

The parameter *β* determines the degree of sparsity, *γ* penalizes excessively large concentrations, and the indicator function, defined as 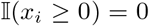 when *x_i_* ≥ 0 and ∞ otherwise, enforces the non-negativity of concentrations.

The optimization problem is to determine the odour **x** most likely to have caused glomerular activations **y**, taking into account both the likelihood and prior. The resulting MAP estimate is given by

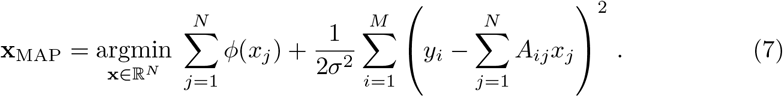

Because the objective function being minimized is strictly convex it has a unique minimum. At this minimum the partial derivative of the objective with respect to each *x_j_* (ignoring for the moment the potential non-differentiabilities introduced in Eq. (6)) is zero, yielding

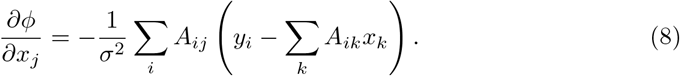

A common approach for solving this equation is to perform gradient descent on the objective function (the right hand side of Eq. (7)) [1], for which the resulting dynamics is

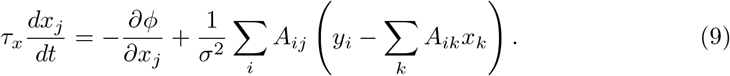

(We recover Eq. (2) if we set *β* and *γ* to zero and *σ*^2^ to 1, and drop that non-negativity constraint in Eq. (6).) These dynamics have a natural interpretation: there’s a leak term due to the gradient of the prior (the first term on the right hand side), feed-forward excitation of the readout unit *x_j_* by the glomeruli (the second term), and recurrent inhibition among the readout units (the third term); see Fig. 1C.

As discussed above, the problem with this approach is that the term 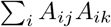 leads to dense connectivity. To remedy that, we can factor out the term in parentheses in Eq. (9) and implement it with a new variable, *λ_i_*, giving us

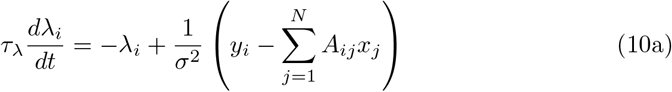

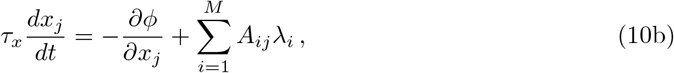

which clearly has the same fixed point as Eq. (9).

The circuitry implied by Eq. (10) is broadly consistent with that of the olfactory system (see Fig. 1D). There is one *λ_i_* for each glomerular input *y_i_*, making it natural to identify *λ_i_* with the activation of a mitral or tufted cell. Mitral and tufted cells likely play different roles in olfactory processing [19], but our theory can be applied to both populations, so, for simplicity, we will refer to them collectively as mitral cells.

Equation (10a) requires each mitral cell to be directly excited by its corresponding glomerulus (*y_i_*) and to be inhibited by the readout units (*x_k_*). Equation (10b) requires the readout units to be directly excited by the mitral cells. This pattern of interaction between the mitral cells and the readout units implies an identification of the readout units *x_j_* with olfactory bulb granule cells, whose main source of excitation is the mitral cells whom they in turn inhibit. Note that in our model the mitral cell/granule connections are symmetric. Since granule cells lack axons, the results of the computation must be read out from the mitral cell activations. This can be done by mirroring the integration of mitral cell input by granule cells as described in Sec. 2.5, ‘Cortical readout,’ below.

### 2.2 Incorporating sister mitral cells

Equation (10a) indicates that each mitral cell *λ_i_* is inhibited by each granule cell *x_j_*, of which there are hundreds of thousands in the mouse [20]. Thus, although the dynamics yield correct inference at convergence, if *A_ij_* is dense we are again faced with implausibly high connectivity. We take inspiration from the olfactory system to show how this problem can be addressed while still performing MAP inference.

So far we have assumed that each glomerulus provides input to one mitral cell (left panel of Fig. 2), but in reality, each vertebrate mitral cell has several dozen ‘sister’ cells (mitral and tufted cells) that all receive input from the same glomerulus [21], (right panel of Fig. 2), and all receive inhibitory feedback from granule cells. This suggests a way to reduce the number of mitral cell/granule cell connections: let each sister mitral cell connect to a different, non-overlapping, set of granule cells. Given that there are at least ~25 sister cells per glomerulus [22,23], that would reduce connectivity by a factor of 25, yielding biologically plausible levels. It turns out that naively implementing such a scheme doesn’t exactly work. However, using this idea, but adding additional cells and connections, this circuitry can indeed perform correct MAP inference.

**Fig 2.**
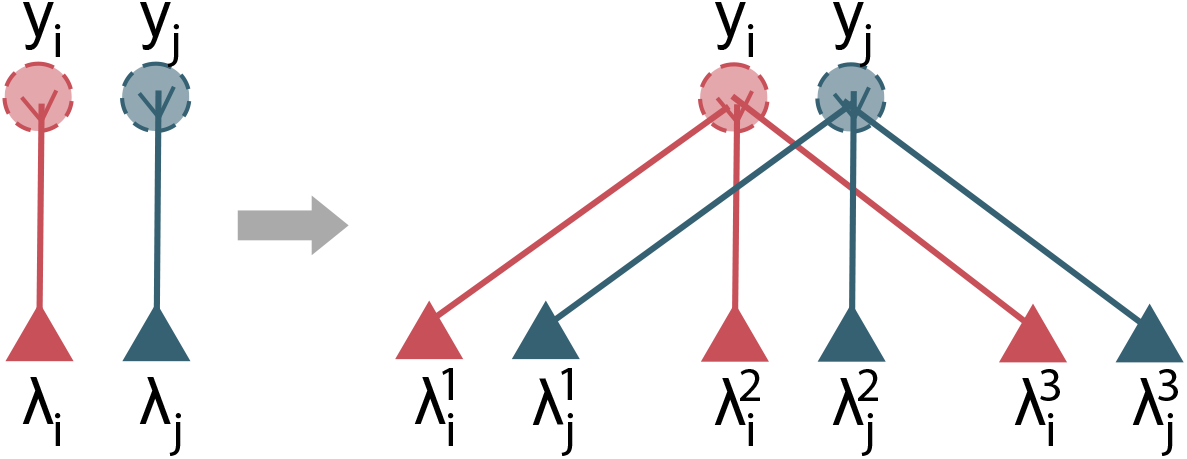
Sister cells. In the vertebrate olfactory bulb, each glomerulus is sampled by not one (left panel) but at least ~25 ‘sister’ mitral and tufted cells [22,23] (right panel, with only three sisters to reduce clutter).

To show that, we start by assuming that the sister cells obey dynamics similar to Eq. (10a),

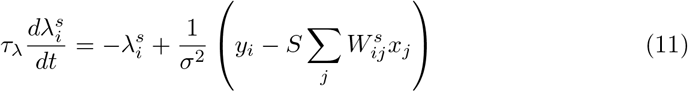

where 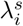 is the *s*^th^ sister cell for glomerulus *i* and *S* is the number of sister cells. Below we will choose 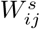 to be zero for all but one *s*, which greatly reduces the number of granule cells that connects to each mitral cell, but for now we leave it arbitrary.

To see how sister mitral cells can perform correct MAP inference, note that the average sister cell activity,

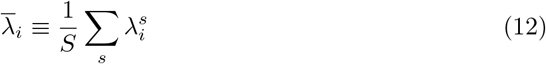

evolves according to

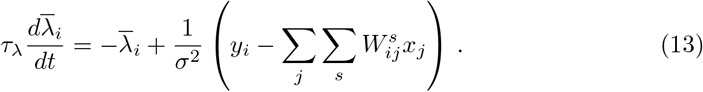

Letting the weights, 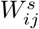, obey

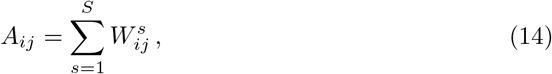

we see that the average sister cell activity evolves according to

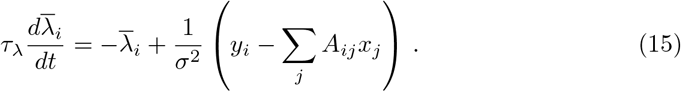

This is identical to Eq. (10a), the time evolution equation for *λ_i_*, implying that under the sister cell dynamics given in Eq. (11), 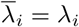. Consequently, rather than computing *λ_i_* from Eq. (10a), we can compute it by simply averaging over the sister mitral cells,

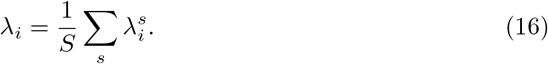

We can, therefore, replace *λ_i_* in Eq. (10b) with the right hand side of Eq. (16), and, so long as the sister mitral cells evolve according to Eq. (11), our model will implement MAP inference.

Equation (14) tells us only that the weights of the sister mitral cells add up to *A_ij_*, but besides that we have complete freedom in choosing them. A trivial choice is 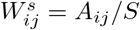 (illustrated in Fig. 3B). However, this doesn’t help, as each sister mitral cell still receives *N* inputs – one from each granule cell. What we want to do instead is make the connections sparse so that, as mentioned above, each sister cell receives input from a different, non-overlapping set of granule cells. If the sets are equal in size, this means each sister cell receives input from *N/S* granule cells, an *S*-fold sparsification (Figs. 3C and D).

**Fig 3.**
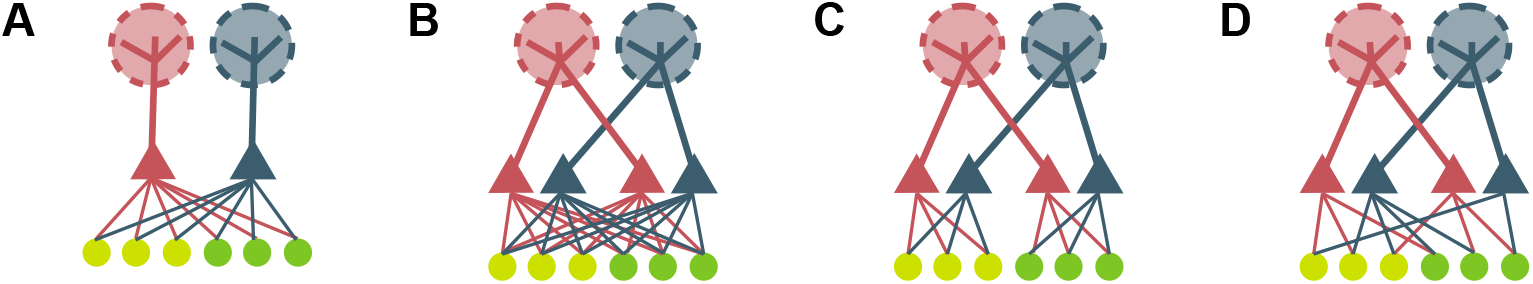
Sparsifying connectivity with sister cells. Each panel shows a schematic of the connectivity between mitral and granule cells. All connections are reciprocal. (A) Original scenario – no sister mitral calls. Connectivity is all-to-all, and the weights are *A_ij_*. (B) A densely connected configuration where every granule cell connects to all sisters on each glomerulus, with weight 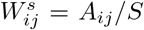. The sister mitral cells recieve the same number of connections as in panel A, but the granule cells recieve twice as many as in panel A (*S* times as as many in general). All-to-all connectivity is thus exacerbated. (C) Blocks of granule cells connect to the same sister from each glomerulus, leading to maximally sparse connectivity. In this example, the first three granule cells connect to the first sister on each glomerulus, and the second three granule cells connect to the second sister; see Eq. (17). Sister mitral cells now interact with a factor of 2 fewer granule cells (*S* fewer in general), sparsifying mitral cell connectivity. (D) A more realistic maximally sparse connectivity pattern. Each granule cell connects to a single, randomly selected sister cell from each glomerulus; see Eq. (18). Sister mitral cells again connect to a factor of *S* fewer granule cells.

We still have a great deal of freedom in choosing the weights. One way would be to first divide granule cells into blocks, and then have the granule cells in each block project to one of the sister cells corresponding to each glomerulus,

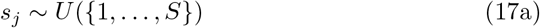

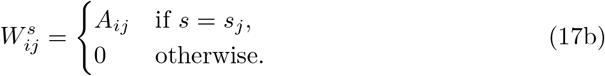

This scheme is shown in Fig. 3C (with the blocks arranged sequentially). More realistic is for each granule cell to connect to a randomly selected sister cell from each glomerulus,

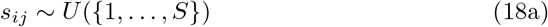

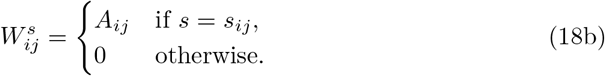

This scheme is shown in Fig. 3D. In either case, each sister mitral cell now receives input from a factor of *S* fewer granule cells. The granule cells still make *M* connections, but, at least in the olfactory system, *M* is relatively small; on the order of 10^2^ – 10^3^.

At this point we have demonstrated that the dynamics in Eqs. (10b) and (11) will lead to MAP inference if *λ_i_* is set to the average of the sister mitral cell activity; that is, if Eq. (16) is satisfied. However, no known cell performs the averaging required in Eq. (16) and then projects to the granule cells, as required in Eq. (10b). We therefore take an alternative strategy: we augment the dynamics to ensure that all the sister mitral cells converge to the average. To do that, we introduce a new cell type (which we will ultimately identify as periglomerular cells) that evolves according to

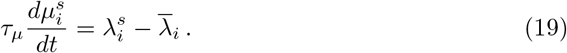

This achieves the desired result: at equilibrium, when 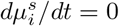, all sister mitral cells associated with glomerulus *i* have the same value – the average sister mitral cell activity. To ensure that 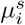 converges to an equilibrium, rather than increasing or decreasing linearly with time, we need 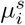 to couple to the sister mitral cells. The natural coupling is linear negative feedback, transforming Eq. (10a) to

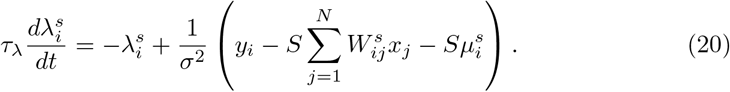

This certainly has the right flavor: positive 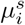 tends to decrease 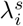, and vice versa, suggesting that all the sister mitral cells will eventually have the same value. But will they have the right value – the value implied by Eq. (10a)? To answer that, we combine Eq. (14) with Eq. (20) to write

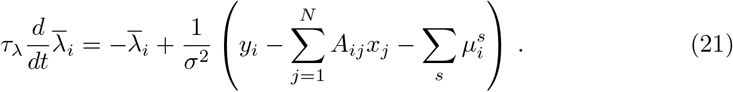

Except for the last term in parentheses, Eq. (21) is exactly the same as the equation for *λ_i_*, Eq. (10a). Note, however, that

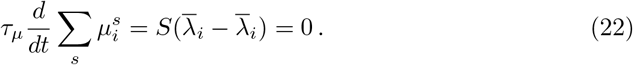

Hence, if we initialize 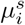 so that 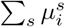 is zero, it will remain zero for all time. In that case, the equation for the sister cell average, Eq. (21), is identical to Eq. (10a). Consequently, each of the sister cells converge to the correct mean, and so we can replace Eq. (10b) with

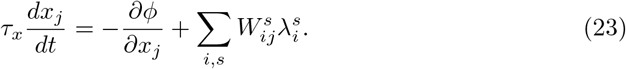

Thus, under the dynamics given in Eqs. (19), (20) and (23), with 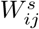 obeying Eq. (14), the network performs MAP inference.

The variables 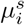 in Eq. (19) are driven by a weighted average of sister cell activations. The observed backpropagation of mitral cell action potentials to the glomeruli [24,25] and the electrical coupling of sisters at the glomeruli [26] might contribute to the neural implementation of just such an average. Thus we have provisionally identified the 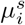 variables with olfactory bulb periglomerular cells because they inhibit the mitral cells and are in turn excited by them [27], and do not receive direct receptor input themselves, reminiscent of the Type II periglomerular cells of Kosaka and Kosaka [28].

### 2.3 Leaky periglomerular cells

The dynamics in Eq. (19) implies that the periglomerular cells, 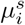, do not leak; i.e., they are perfect integrators. This is at odds with biology, since we imagine that integration is performed by neuronal membranes, and neuronal membranes are leaky [29] – though periglomerular cells may be less leaky than most other neurons because of their very high input resistance [30]. We can introduce a leak term into the dynamics,

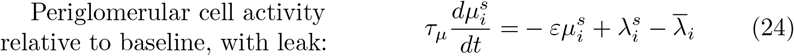

where *ε* sets the magnitude of the leak. One advantage of introducing this leak is that we no longer have to worry about initializing the 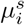 to have zero mean, since with a leak term the mean periglomerular activation decays to zero,

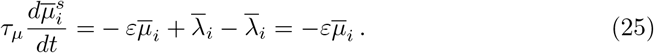

The price we pay is that the system no longer computes the MAP solution exactly. As we show in Methods, Sec. 4.1, the system of equations minimize the wrong objective,

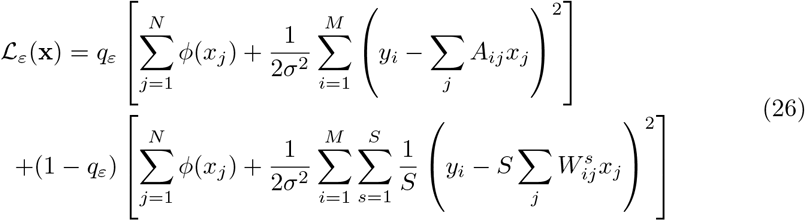

where

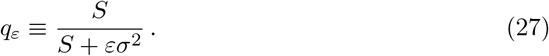

In the limit of no leak (*ε* → 0, so that *q_ε_* → 1), we recover the correct objective (compare to Eq. (7)). For non-zero leak, the objective differs from the MAP objective so solutions are likely to differ from the MAP solution. However, as we show numerically in Figs. 9 and 10, for biologically relevant values of *ε* these deviations are small.

Note that as the number of sister mitral cells S increases, *q_ε_* approaches 1. Naively, this suggest that we should recover the true objective in the large *S* limit. However, this naive expectation ignores the fact that there is a factor of *S* in the second term in Eq. (26); in the large *S* limit, this cancels the 1/*S* dependence in (1 – *q_ε_*). Consequently, it is not immediately clear how inference depends on *S* when there is nonzero leak. We addressed this numerically, and found that the error relative to the MAP solution increases monotonically with the number of sisters; see Fig. 10.

### 2.4 Implementation in neural circuitry

The mitral cell dynamics in Eq. (20) and those for the periglomerular cells in (19) are already in a form suitable for implementation by neural circuitry. However, the granule cell dynamics in Eq. (23) cannot be implemented directly because of the presence of not-everywhere-differentiable terms in the prior, *ϕ* (see Eq. (6)). We thus implement related, neurally plausible, dynamics that has the same fixed point. Specifically, we note that at the fixed point of Eq. (23), *x_j_* satisfies

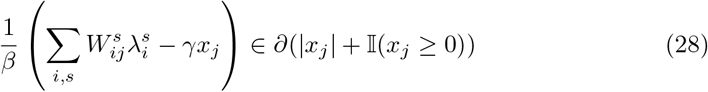

where *∂* is the subgradient operator [31]. If at the solution *x_j_* > 0, then the subgradient operator reduces to the ordinary gradient and yields the value 1, and we have

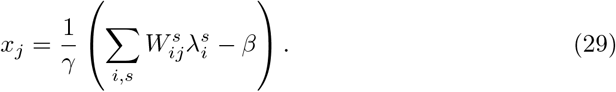

On the other hand, when *x_j_* = 0 the subgradient is the set (-∞, 1], so we have

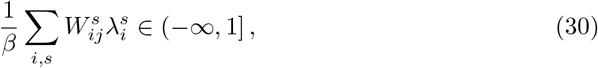

which we can write as an inequality,

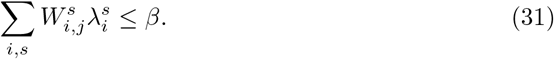

Thus, *x_j_* = 0 whenever Eq. (30) is satisfied; combining that with Eq. (29) (which is valid when *x_j_* > 0), we have

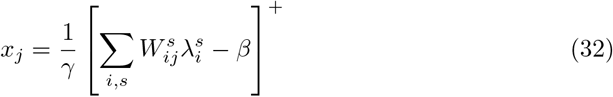

where [.]^+^ is the threshold-linear operator. A neural implementation of this function would have Xj responding instantaneously to changes in the mitral cell activations 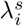, which is implausible. Instead we employ a membrane voltage variable *v_j_* which integrates the mitral cell input and interpret *x_j_* as the resulting firing rate. The full set of equations describing the model is

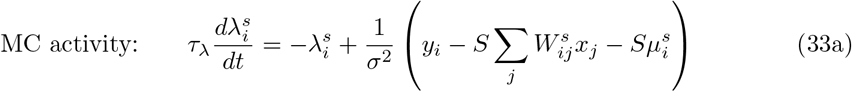

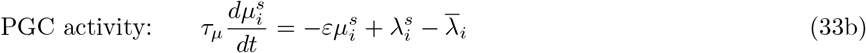

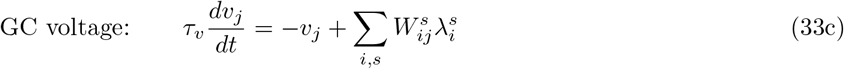

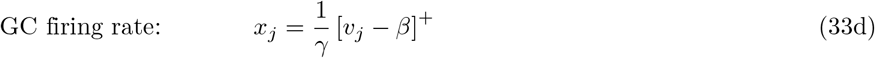

where the weights 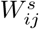 satisfy Eq. (14). Equations (33a) and (33b) correspond to Eqs. (20) and (19), respectively, and Eqs. (33c) and (33d) implement Eq. (32).

A circuit that implements these equations is shown schematically in Fig. 4. As promised, each mitral cell interacts with only a subset of the granule cells, as in Fig. 3D. This reduces mitral-granule connectivity by a factor of *S* (though the *total* number of mitral-granule synapses stays the same due to the introduction *S* sister mitral cells per glomerulus). The information from the other granule cells is delivered indirectly to each mitral cell via the influences of the glomerular average of the sister cell activations and periglomerular inhibition.

**Fig 4.**
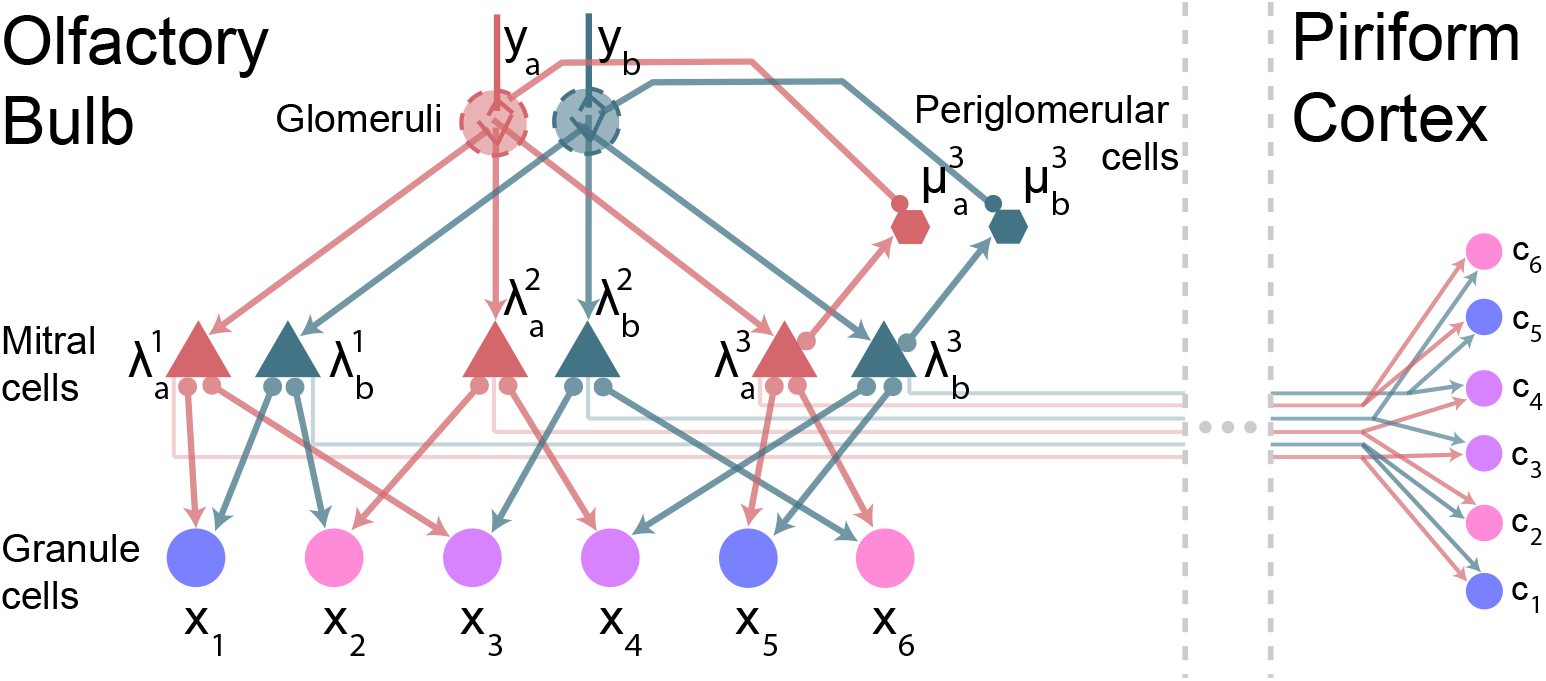
Sparsely connected inference circuit, with readout in piriform cortex. Arrows and filled circles indicate excitatory and inhibitory connections, respectively. Each sister cell 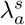 receives input *y_a_* from a glomerulus and interacts with a subset of the granule cells *x_j_*, reducing connectivity by a factor of *S* (*S* = 3 in this example). Information is shared between granule cells through the periglomerular cells 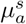 (only ones with *s* = 3 are shown, for clarity). Those cells receive excitatory input from the mitral cells and inhibitory input corresponding to the average sister cell activity (as required by Eq. (33b)). The inhibitory input comes from backpropagating action potentials, which travel along mitral cell apical dendrites and are available at the glomeruli [24, 25]. The result of the inference is simultaneously available in granule cells and neurons in piriform cortex (see Sec. 2.5, ‘Cortical readout’.)

A strong prediction of the above implementation of our model is that when the periglomerular cells are perfect integrators (*ε* = 0 in Eq. (33b)) all sister cells associated with any one glomerulus converge to the same value for each odour. It turns out, though, that convergence to the same value is not a general feature of our model; it’s a consequence of the specific choices we made. To see that, we can simply make the linear change variables,

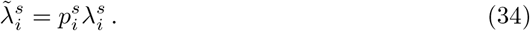

This has very little effect on the equations because this change of variables in Eq. (33) simply introduces a multiplicative scaling in several terms in that equation. It does, though, have a quantitative effect: sister mitral cells, 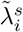, no longer converge to the same value. Instead, any two sister cells converge to the same, odour-independent, ratio,

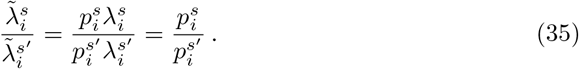

Thus, our model makes the very general prediction that, in steady state, the activity of any two sister mitral cells sharing the same glomerulus will have the same ratio, independent of odour.

This prediction holds, however, only under a linear change of variables. Under a nonlinear change of variables, 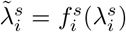, the ratio of sister mitral cell activity is given by

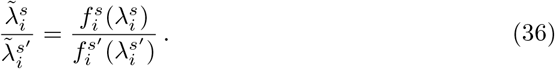

At convergence, this will depend on 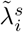 and 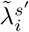, and thus on odour. This changes our prediction to: if periglomerular cells are perfect integrators, there is a nonlinear transformation of the mitral cell activity such that at convergence, the ratio of transformed sister mitral cell activity is independent of odour. While more difficult to test than the simpler prediction above, with sufficient data it should be possible.

Finally, the presence of a leak also modifies this prediction. When *ε* > 0, Eq. (33b) tells us that, at convergence,

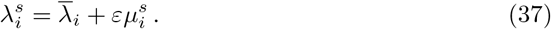

Consequently, the ratio of sister mitral cells will acquire an odour dependence. However, in our simulations, that dependence was weak (see Fig. 10C).

### 2.5 Cortical readout

In our model, odour concentration is stored in granule cells, which don’t project outside of the olfactory bulb, and in fact lack axons entirely [27]. Thus, the granule cells can’t communicate any information to the rest of the brain. This can be remedied by projecting mitral cell activity to cortical readouts 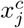 via projection weights 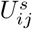

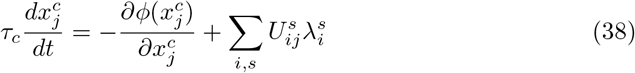

In this circuit each cortical neuron 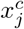 is excited by the sister cells in the same way as the granule cells in the bulb, but is not required to provide feedback to the bulb. When computation in the bulb converges, we have 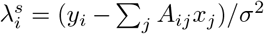 (see Eq. (33a) and recall that sister mitral cells all converge to the same activity), so that

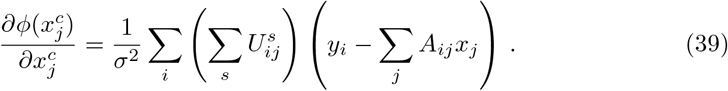

Thus as long as the projection weights to the cortex satisfy

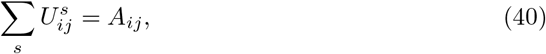

(analogous to Eq. (14)) then cortical neuron 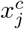 will have the same fixed point as granule cell *x_j_*. This means the output of the computation in the bulb can be read out in the cortex via a 1-to-1 correspondence between granule cells and cortical neurons. Thus basic olfactory inference can be performed entirely within the bulb, with the concomitant increase in computational speed, and the results can be read out in the cortex. As cortical feedback to the bulb, in particular to the granule cells, does in fact exist [27], its role may be to incorporate higher level cognitive information and task contingencies into the inference. We leave the exploration of these ideas to future work.

### 2.6 Simulations and linear analysis

To investigate the behaviour of the system, we performed a series of simulations using the model summarized in Eq. (33). The three questions that guided our choice of simulations, along with brief answer, are:

1. What do the dynamics of sister cells look like? Our analysis in the main text shows only that if the dynamics converges, it yields the MAP solution. However, we have not shown that the dynamics necessarily converges, or said anything about transient behaviour. Thus the first goal of our simulations was to check convergence empirically, and to qualitatively assess the transient dynamics and its biological plausibility. Answer: we show numerically that the dynamics do indeed converge, and show analytically that solutions are stable for the parameter regime of interest.
2. What is the effect of non-zero periglomerluar leak (*ε* > 0)? The dynamics in Eq. (33) yields the MAP solution at convergence only when the periglomerular cells have zero leak, yet any biological implementation of these dynamics will have non-zero leak. It is important to determine the extent to which realistic values of the leak affect the dynamics and the inference solutions. Answer: for realistic values of the leak, the effect on MAP inference is small.
3. Finally, how do the various parameters affect the transient dynamics, and which parameters are most important? In particular, does the dynamics become qualitatively unrealistic when some parameters are changed within biologically reasonable ranges? Answer: the transient dynamics is extremely robust to parameters, and exhibits very little change over a broad range.

Below we expand on these answers.

#### 2.6.1 System dynamics with sister cells

In our simulations we used the base parameters given in Table 1 below; departures from those parameters will be flagged. Figure 5 shows typical activity patterns for mitral, periglomerular and granule cells. Panel A shows the response of all four sister mitral cells from a representative glomerulus, and panel B shows the response of the corresponding periglomerular cells. Although there is some initial variability in the responses - in particular decaying oscillations – the sister cells converge to the same value, as expected. Panel C shows granule cell activity, and demonstrates that the readout converges to the MAP solution within a few hundred milliseconds. This is reflected in the root mean square (RMS) error between the granule cell responses and the MAP estimate, which decays exponentially (panel D).

**Table 1.** Base parameters used in the simulations.

| Parameter | Description | Base Value |
| --- | --- | --- |
| $M$ | Number of glomeruli | 50 |
| $N$ | Number of granule cells | 1200 |
| $S$ | Number of sister cells per glomerulus | 4 |
| $n$ | Number of components present in the true odour | 3 |
| $\sigma^2$ | Receptor variance | $10^{-2}$ |
| $\beta$ | $\ell_1$ penalty | 3 |
| $\gamma$ | $\ell_2$ penalty | 1 |
| $\tau_v$ | Granule cell membrane time constant | 35 ms |
| $\tau_\lambda$ | Mitral cell membrane time constant | 50 ms |
| $\tau_\mu$ | Periglomerular cell membrane time constant | 35 ms |
| $\varepsilon$ | Periglomerular cell leak | 0 |
| $A_{ij}$ | Elements of the affinity matrix | $\mathcal{N}(0, 1/M)$ |

**Fig 5.**
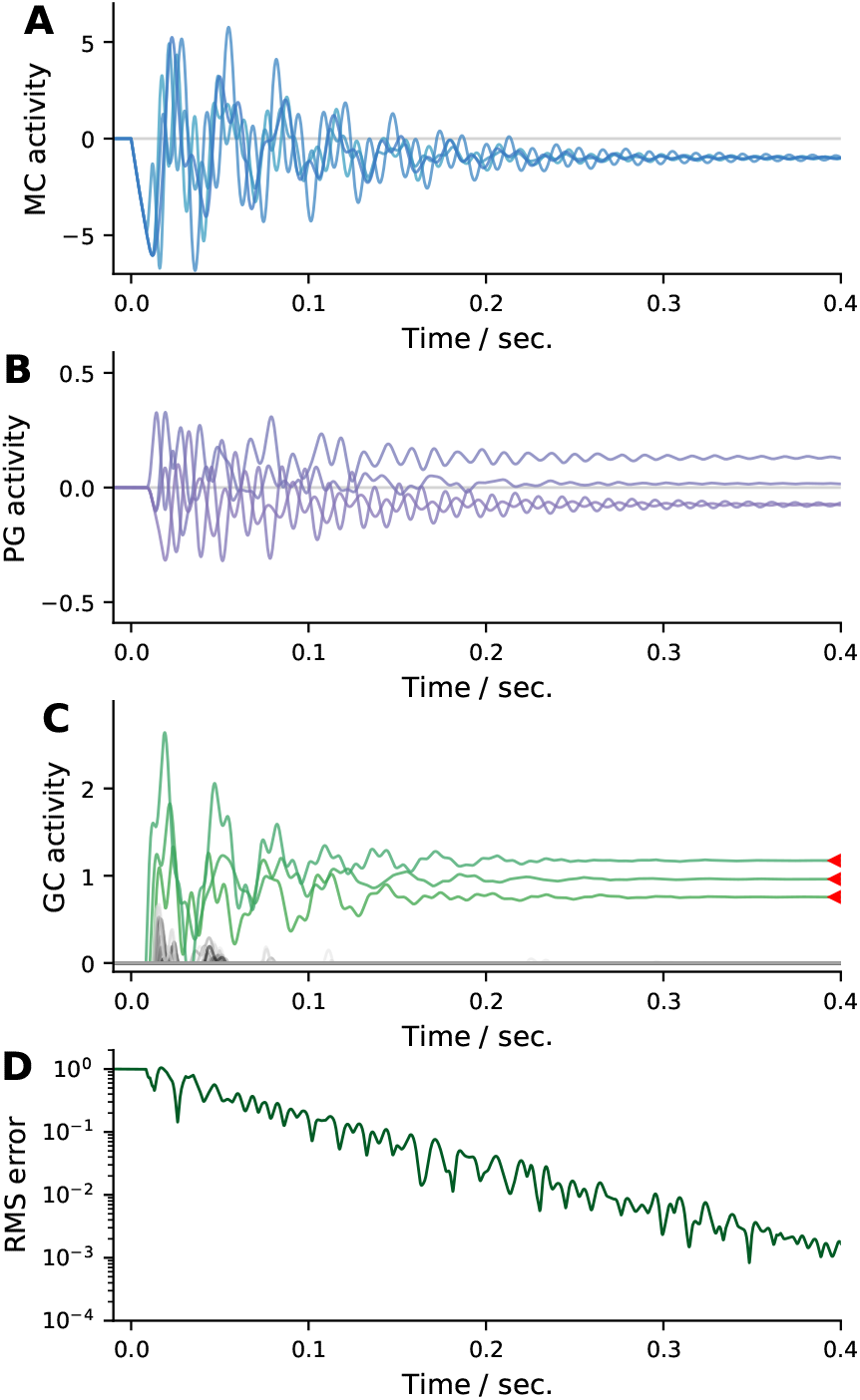
Example dynamics for a network with base parameters (Table 1). (A) All four sister mitral cells from a representative glomerulus. Although the sister cells start off with identical activations, their activities quickly diverge due to their differing connectivities to the granule cells. Ultimately, however, they converge to the same value. (B) Periglomerular cells from the glomerulus in panel A. (C) Granule cells. Red arrows indicate the MAP solution for the three presented odours. Granule cells representing these odours are shown in green, the others in gray. After an initial period of activity, the system settles into the MAP solution. (D) Time course of the root-mean-square error between the granule cell activations and the MAP solution normalized to its initial value, indicating convergence to the MAP solution.

In Figure 6 we show typical dynamics when there are *S* = 1, 8 and 25 mitral cells per glomerulus. In all cases, the granule cells converge to the MAP solution within a few hundred milliseconds. The main difference between the three values of *S* is that when *S* = 1, convergence to the MAP solution is slightly faster than when *S* > 1, as indicated by the slightly steeper RMS error cuves in the bottom left panel. Otherwise, the dynamics in all three cases is qualitatively similar.

**Fig 6.**
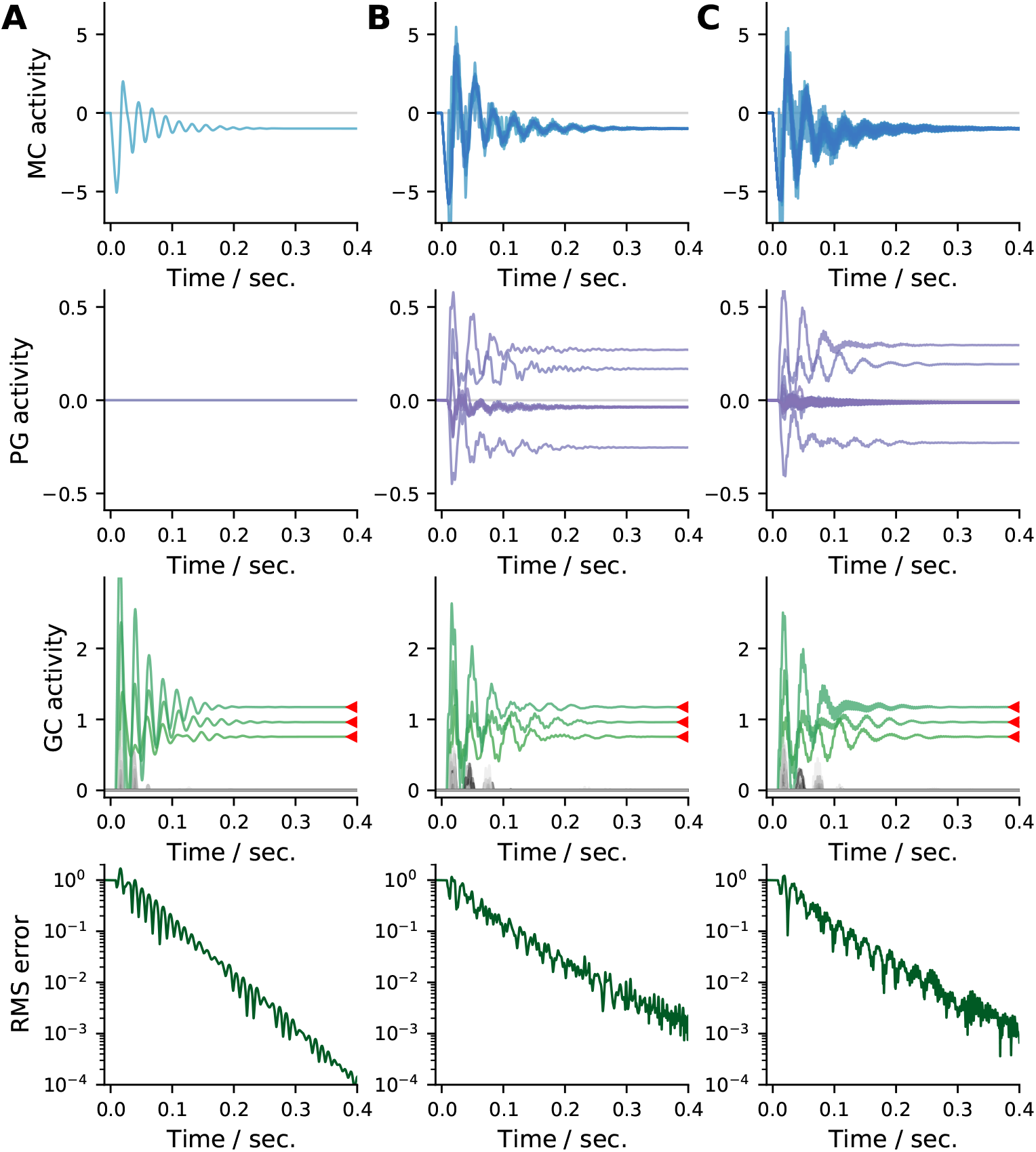
Typical dynamics as the number of mitral cells per glomerulus, *S*, is varied relative to the base model given in Table 1. (A) *S* = 1, (B) *S* = 8, and (C) *S* = 25 mitral cells per glomerulus, compared to *S* = 4 in Fig. 5. Note the lack of periglomerular cell activity and slightly faster convergence for *S* = 1. Otherwise, the dynamics is qualitatively similar for all values of *S*, including convergence to the MAP solution.

##### Linear analysis

Given that our network was designed to perform MAP inference, the asymptotic behavior shown in Fig. 5 and Fig. 6 is relatively unsurprising. However, our analysis so far says nothing about the transient behavior, which, as can be seen in Figs. 5 and 6, is characterized by large oscillations. To understand this behavior – in particular how stability and oscillation frequency depend on the parameters, including the number of sister mitral cells – we need to analyze the dynamics.

Because of the rectifying nonlinearity, that is hard to do exactly. However, our simulations so far (in particular the granule cell activity in Figs. 5 and 6) suggest that the composition of the ‘active’ set of granule cells stabilizes before the dynamics terminates. Once this active set has stabilized, so that the rectifications remain within the linear regime, the granule cell activations *x_j_* can be replaced by their corresponding voltage variables *v_j_*, and the system becomes linear. We thus performed linear analysis of Eq. (33) around a solution with *n* active granule cells, and used the results to both explain the transient behavior we have seen so far, and guide further investigation of the system.

The linearized dynamics relative to their input-dependent fixed-point for *n* active granule cells is described by the equations (Methods, Sec. 4.2)

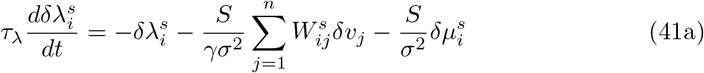

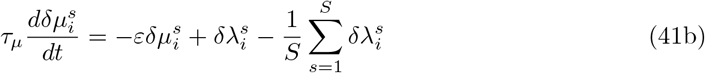

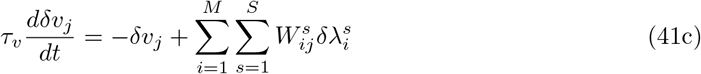

where the *δ* in front of each variable indicates a small deviation from the fixed point predicted by MAP inference. Note that we replaced the granule cell activations *x_j_* by their membrane voltages *v_j_*, as motivated above, and that the indexing of the latter variables is over the active set of *n* granule cells.

To solve these equations, we let the dynamical variables have the time dependence *e^ξt^*, which results in an eigenvalue equation for *ξ*. That equation can’t be solved exactly (at least not for all eigenvalues), but the approximate eigenvalues, derived in Methods, Sec. 4.2, are reasonably close to the true ones, as shown in Fig. 7. In that figure, eigenvalues near blue markers are for modes involving only mitral and periglomerular cells (*δv_j_* = 0), while eigenvalues near orange and red markers are for modes involving granule cells as well (*δv_j_* ≠ 0).

**Fig 7.**
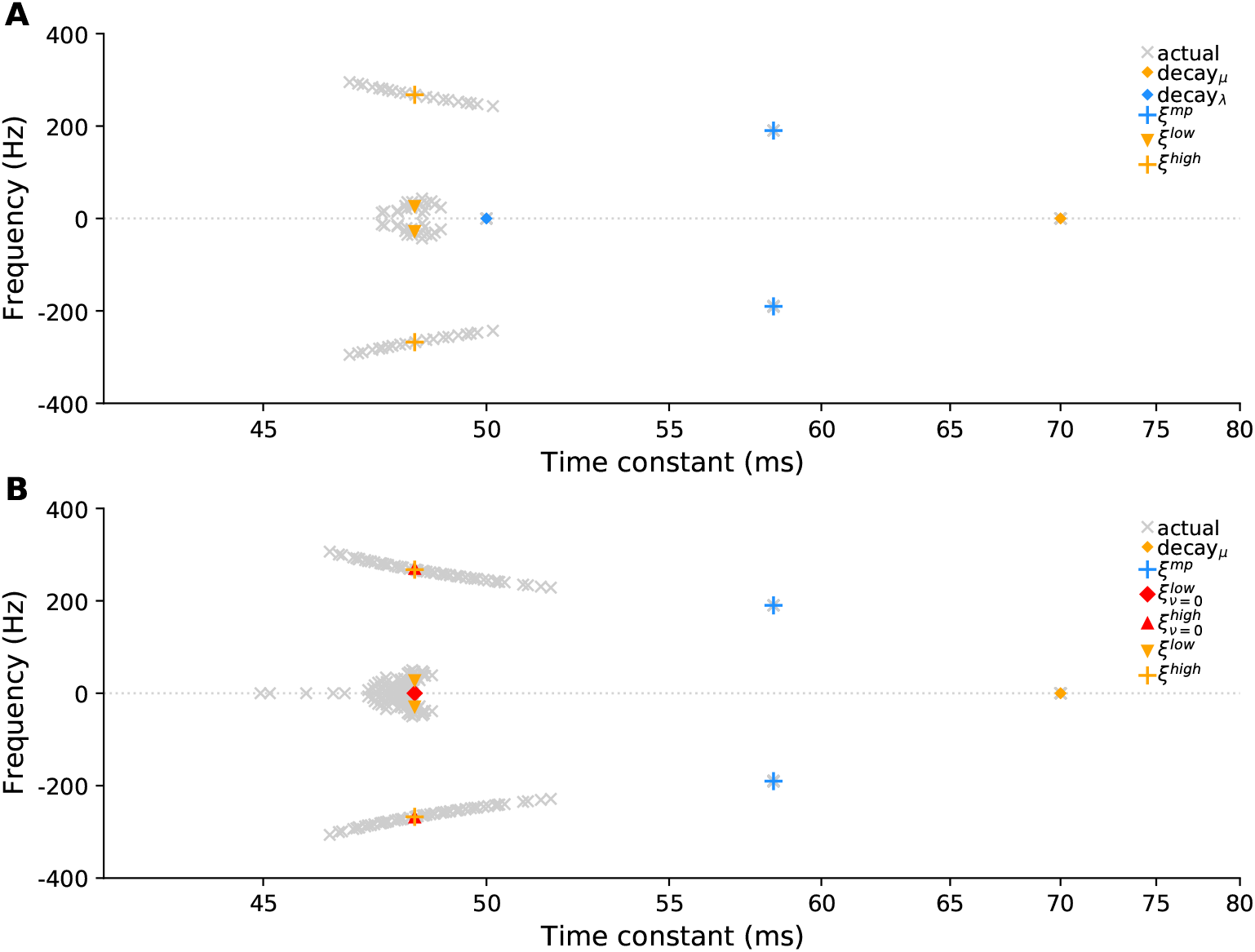
Eigenvalues of the linearized system computed numerically (gray crosses) and analytically (colored symbols). Eigenvalues labeled ‘decay_*μ*_’, Eq. (64), ‘decay_*λ*_’, Eq. (65) and ‘*ξ^mp^*’, Eq. (66), are exact; those with the superscripts ‘low’ or ‘high’, Eq. (83), are approximate. (A) *n* = 20 active granule cells. (B) *n* = 55, so that *n* > *M*. In both panels, blue eigenvalues are for modes involving only mitral and periglomerular cells, modes for orange and red eigenvalues also involve granule cells. The time constant (*x*-axis) and frequency (*y*-axis) for each eigenvalue *ξ* are –1/Re(*ξ*) and Im(*ξ*)/2*π*, respectively. We used large *n* to better show the spread of eigenvalues. Parameters from Table 1, except *S* = 25 and *ε* = 1/2, and *n* depends on the panel. See Sec. 4.2 for additional details.

Each gray cross in Fig. 7 corresponds to a mode of the system, and near the equilibrium the activity consists of a sum of these modes. However, because the modes cluster, the system admits only a handful of behaviors, which we summarize as follows:

1. Low frequency oscillations, ‘*ξ^low^*’, whose frequency does not change with added sisters (see Methods, Eq. (87));
2. Two high frequency oscillations, ‘*ξ^high^*’, which involves all cell types, and ‘*ξ^mp^*’, which involves only the mitral and periglomerular cells. The frequency of these oscillations increases with added sisters (see Methods, Eq. (85));
3. Purely decaying modes (no oscillations), ‘decay_*μ*_’, (orange diamond in Fig. 7), which has a decay rate that is proportional to *ε*, and ‘decay_*λ*_’, which is present only when *n* < *M* (blue diamond in the top panel of Fig. 7). The latter involves the mitral and periglomerular cells, but not the granule cells.

Figure 7 shows the eigenvalue spectrum for only one set of parameters. What about other choices? The time constants, *τ_λ_, τ_μ_*/*ϵ* and *τ_v_*, set the time scale for decay. The other relevant parameters are the number of sister mitral cells, *S*, and *γ* and *σ*^2^; the latter two appearing in Eq. (41a). As we show in Methods, Sec. 4.2, these have two main effects. First, the oscillation frequencies *ξ^high^* and *ξ^mp^* scale as 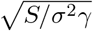 (Eq. (85)) and 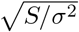 Eq. (66), respectively, as shown in Fig. 8A and B. The second effect is on the decay time constants, which are mainly independent of *S*, except when *S* = 1; in that case there is no *ξ^mp^* mode, which can affect the decay rates (see Fig. 8C). For a detailed analysis of the effect of parameters on the transient behavior, see Methods Sec. 4.2.

**Fig 8.**
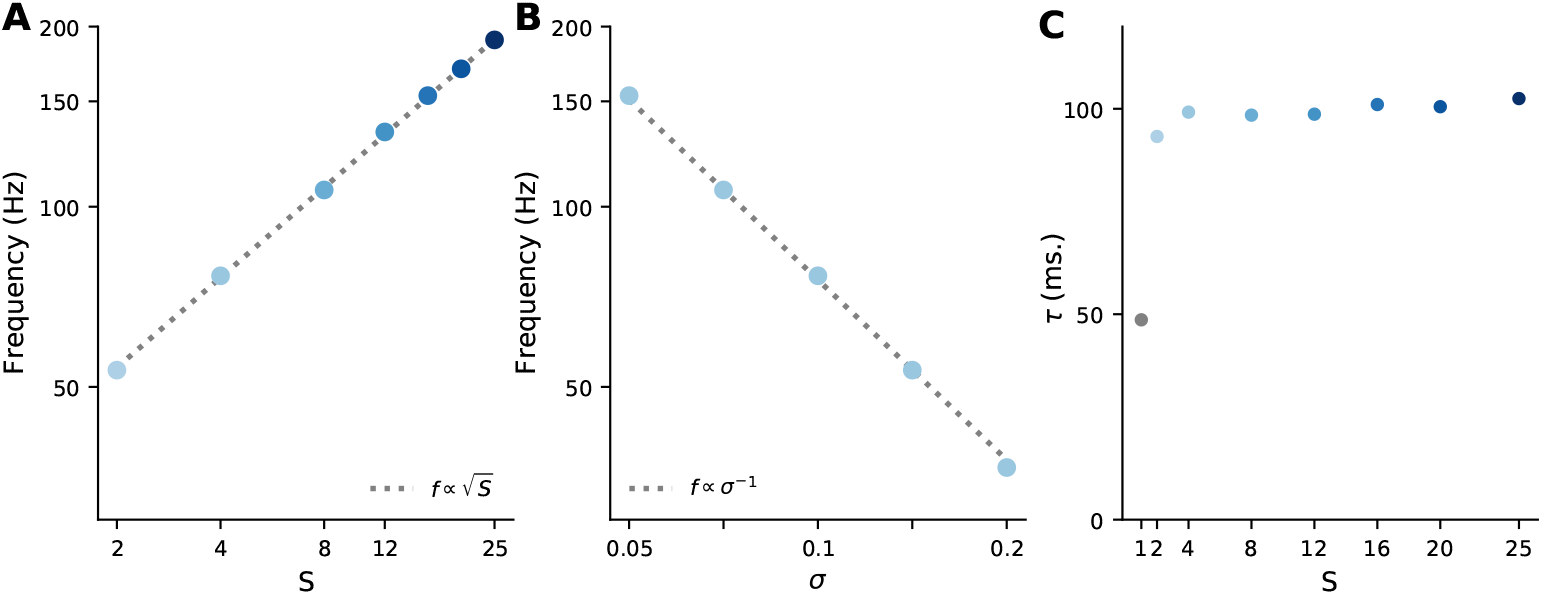
Transient properties of inference dynamics. (A) Dependence of the high-frequency mode of the mitral cell responses on the number of sisters, *S*. (B) Dependence on the standard deviation of the noise, *σ*. In both panels, the dashed lines are fits predicted by the linear analysis. (C) Dependence of the mitral cell decay time constant on the number of sisters *S*. Going from *S* = 1 to *S* = 2 sisters per glomerulus decreases the decay time constant by a factor of about 2. A large change in decay rate from *S* = 1 to 2, followed by a slower change for *S* ≥ 2, is typical, although the details are parameter-dependent. Other parameters as in Table 1.

The linear analysis can also tell us whether the MAP equilibrium can be unstable. We show in Methods, Sec. 4.2.2 that it is stable in the parameter regime of interest, which is *σ*^2^ ≪ 1; we did not investigate stability when *σ*^2^ isn’t especially small.

#### 2.6.2 The effect of periglomerular leak

As discussed in Sec. 2.3, our network performs MAP inference only if the periglomerular leak, *ε*, is zero. What happens in the realistic case, when it’s not zero? Here we address that question through simulations. From Eq. (33b), we see that the effective time constant of the periglomerular cells is *τ_μ_*/*ε* = 35 ms/*ε* (see Table 1), so relevant values of *ε* are near 1.

In Fig. 9 we show typical dynamics for *ε* = 1 and *ε* = 2. Non-zero values of the leak mean the system no longer performs MAP inference; that’s reflected in the plateauing of RMS error relative to the MAP solution. In both cases however, the effect on granule cell activity is small. Non-zero leak also means that the periglomerular cells are no longer able to force sister cells to the same value at convergence. This is increasingly visible as the leak increases (compare granule cell activity at convergence in panels A and B of Fig. 9). Again, though, the effect is small.

**Fig 9.**
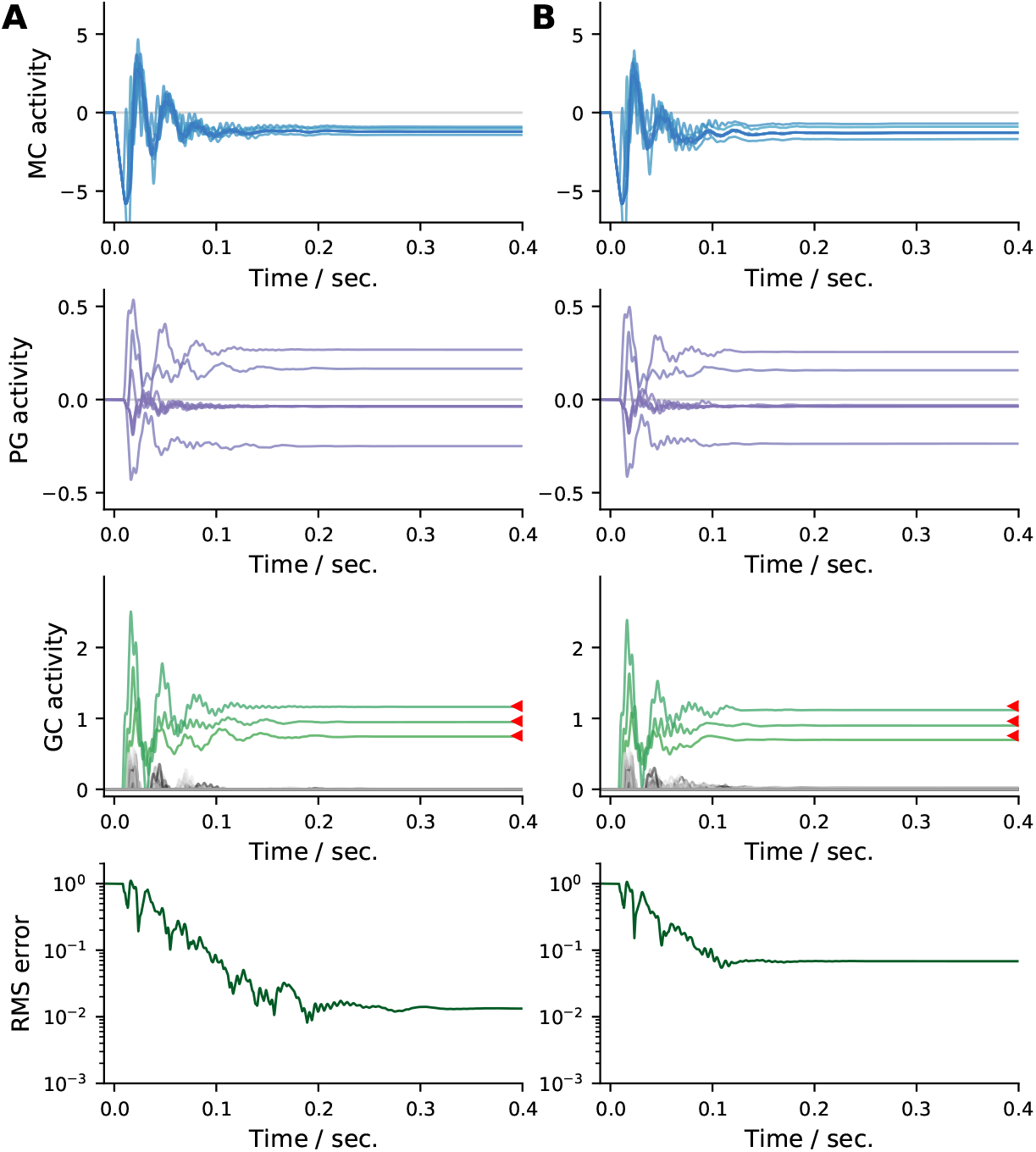
Effect of periglomerular leak *ε* on the dynamics. Parameters from Table 1, but with the number of sisters S fixed at 8 and the leak at (A) *ε* = 1 and (B) *ε* = 2. Note that sister mitral cells no longer converge to the same value (top row). Increasing the leak results in higher RMS error relative to the MAP estimate (red triangles).

As discussed in Sec. 2.3, the effect of the periglomerular leak depends on the number of sister mitral cells, *S*. In Fig. 10A we plot the asymptotic RMS error versus *ε* as we very *S*. Increasing the number of sisters increases the steady state RMS error relative to the MAP solution, but past about *S* = 8, the number of sisters has very little effect on the error. In Fig. 10B we plot the spread in sister cell activity relative to the mean for *S* = 25, showing that it remains small for large values of the leak.

**Fig 10.**
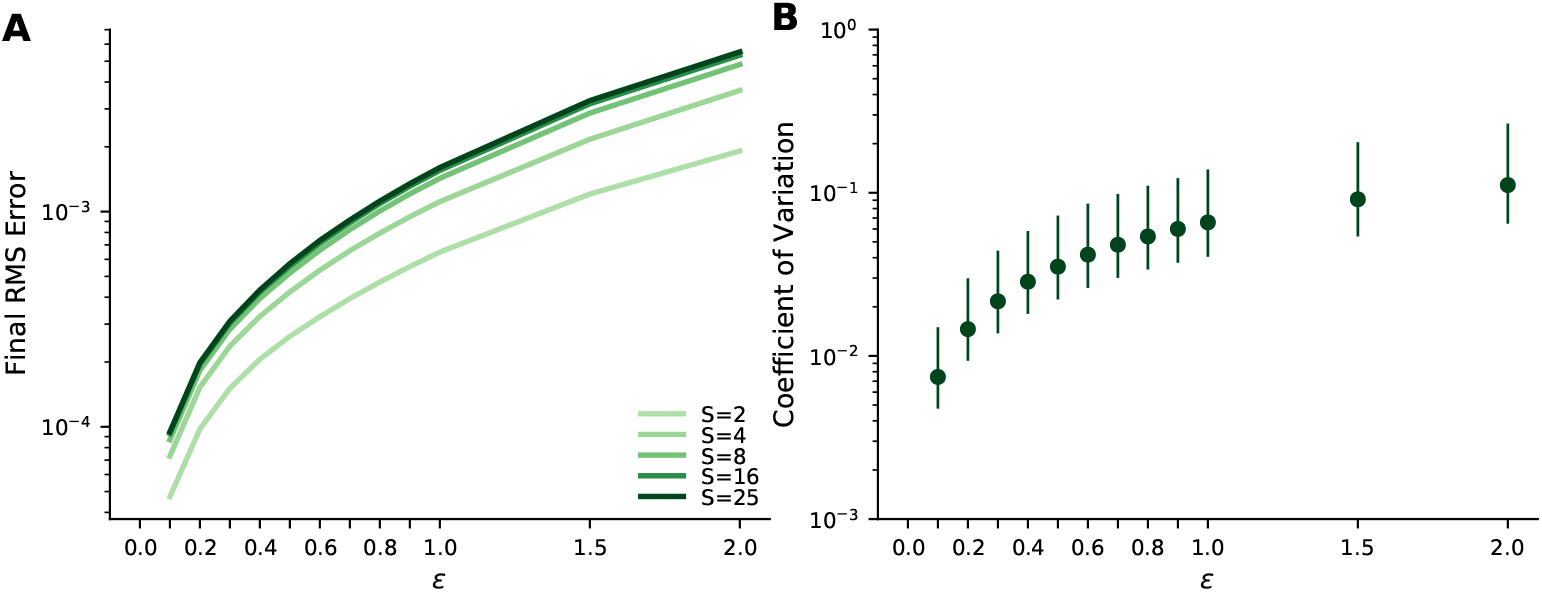
Properties at convergence versus periglomerular leak, *ε*. (A) Mean RMS error at convergence as a function of leak for different numbers of sister cells per glomerulus. The RMS error increases with leak and with the number of sister cells, but never gets very large. Means are computed over 10 different odours and 5 different olfactory bulbs. (B) Median (dots) and inter-quartile range (lines) of the coefficient of variation (CV; standard deviation divided by the mean) of the ratios of sister cell activity at convergence as a function of the leak parameter *ε*. CVs are computed over the ratios of all unique pairs of 25 sisters in a glomerulus, percentiles are computed over all glomeruli in response to 10 different odours across 5 different olfactory bulbs. Median CVs remain low even for high values of leak. Parameters from Table 1, except for *S* and *ε*.

#### 2.6.3 Robustness

Finally, we examined how the parameters affect the dynamics in the non-leaky setting. Because the system always arrives at the MAP solution, we focus on the transient dynamics, and in particular on whether the system remains within a biological plausible range.

##### Number of odours, *n*

Intuitively, we expect that as the number of odours, *n*, increases, the inference problem will become harder. In Fig. 11A we show an example with 10 odours present (dashed gray lines), and indeed we see that inference (blue lines) is not great: although the odours that are present are correctly inferred, many odours that are not present are inferred as well. Figure 11B corroborates this: the number of recovered odours exceeds the number of true ones, n, when n is above about 3. However, for all values of n tested, the dynamics converges to the MAP solution at similar rates, as shown in Fig. 11C.

**Fig 11.**
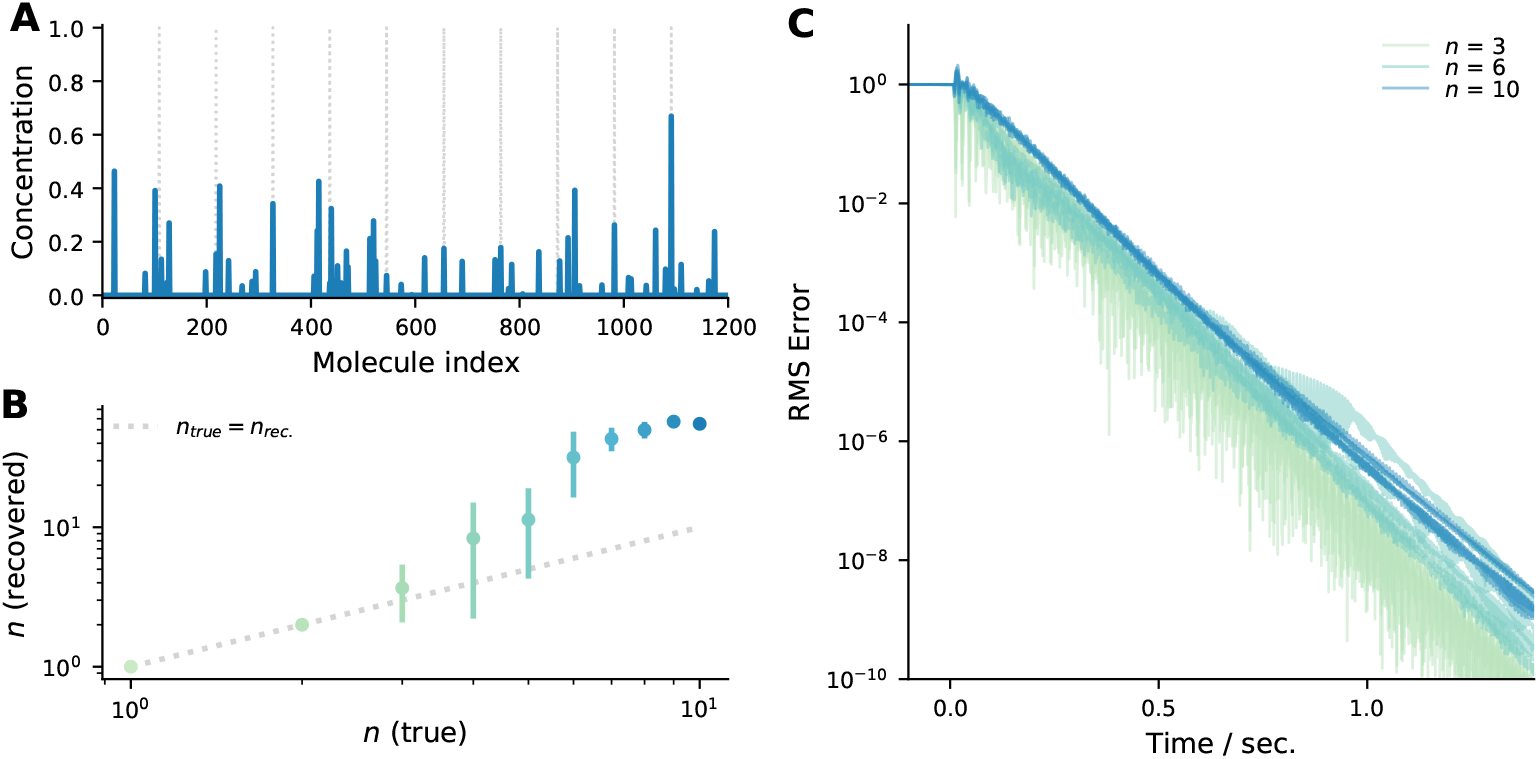
The effect of the number of odours, *n*, on inference. (A) The MAP estimate for an odour with *n* =10 components contains many more than 10 non-zero values. (B) Number of odours inferred using MAP inference versus the true number of odours, *n*. Above about *n* = 3, extra odours are inferred. An odour was considered inferred when granule cell activation was greater than 10^−2^. (C) Time course of the RMS error between the granule cell activations and the MAP estimate, for 6 different trials at each *n*. Parameters from Table 1, except *S* = 8.

##### Number of cells

So far our simulations have used *M* = 50 glomeruli and *N* = 1200 granule cells. However, most olfactory systems are much larger than this (we used smaller populations solely to speed up simulations). For example, in the fly *M* ≈ 50 and *N* ≈ 2500, while in the mouse, *M* ≈ 1000 and *N* ≈ 10^6^. In Fig. 12 we show circuit dynamics for larger systems; up to M = 200 and N = 4800. Consistent with the fact that the linear analysis (Methods, Sec. 4.2) doesn’t predict a strong dependence on size, the dynamics are qualitatively similar to our simulations with small *M* and *N*, and the MAP solution is achieved within a similar time frame.

**Fig 12.**
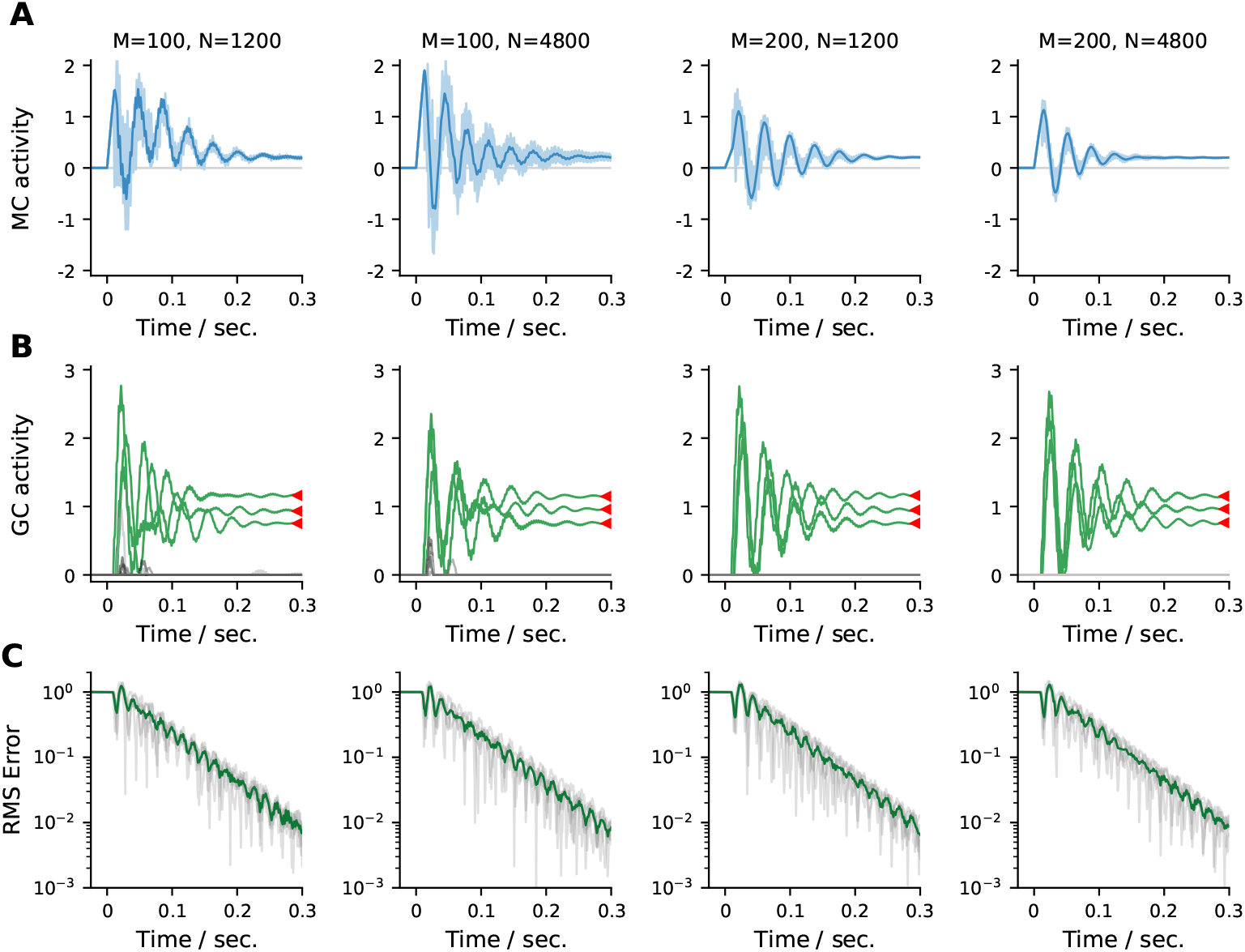
Dynamics for different numbers of glomeruli, *M*, and granule cells, *N* (specified at the top of each column), are qualitatively similar. (A) Mitral cells. The activity of one mitral cell is highlighted with a dark trace; the remaining sisters are overlayed with a light trace. (B) Granule cells. Granule cells representing molecules present in the true odour are colored green and the 10 most active granule cells representing molecules *not* present in the true odour are colored gray. Granule cell dynamics are all qualitatively similar and converge to the MAP solution (red triangles). (C) RMS error between the granule cell activity and the MAP estimate for 5 random systems of the same size as in the previous rows (gray) and the average (green). All systems converge with similar dynamics to the MAP estimate. Remaining parameters are from Table 1, except *S* = 25.

### 2.7 Application to other models

Our model used sister mitral cells to sparsify connectivity in a circuit performing inference under a linear model of olfactory transduction with Gaussian noise. Our approach, however, is quite general, and can be applied to more complex models. For example, in [2] the authors also consider a linear model of olfactory transduction, but with Poisson noise and a spike-and-slab prior on odour concentrations. Equations (28) and (29) from their model translate, with minor redefinitions to be consistent with our notation, and minor simplifications to reduce clutter, to

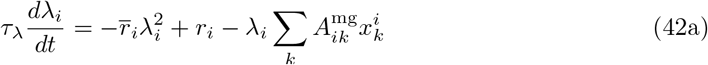

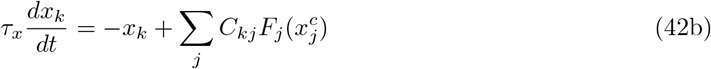

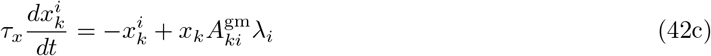

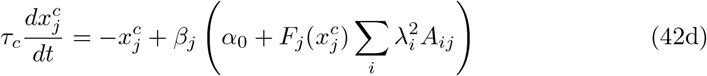

where the connectivity matrices 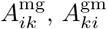 and *C_jk_* are related to *A_ij_* by

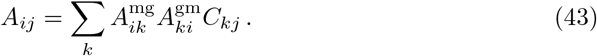

There are several differences between this model and ours. First, the input, which in our model was *y_i_*, is now stochastic: *r_i_* is the number of spikes in a bin of size Δ*t* generated from a Poisson process with rate proportional to *y_i_*, and 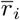 is the expected number of spikes. Second, the granule cells and mitral cells communicate via dendro-dendritic connections at “spines”, denoted 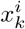; this results in several connection matrices rather than just one. Third, the cortical readout, 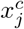, feeds back to the olfactory bulb. Fourth, the nonlinearity, 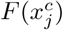, which is defined in terms of the digamma function, is very different from ours. And fifth, the equation for the mitral cells has a term 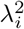 on the right hand side.

What they have in common is that the connectivity matrices, 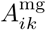 and 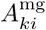, are dense, and so would require mitral cells to interact with nearly all granule cells. This results in the same all-to-all connectivity problem that we highlighted in Eq. (10). But it can again be fixed using sister mitral cells and periglomerular cells,

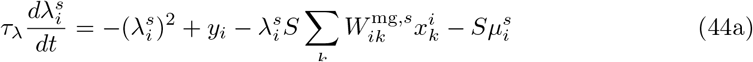

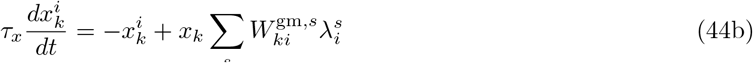

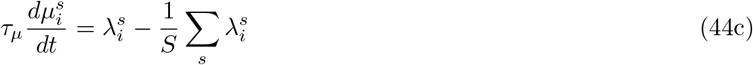

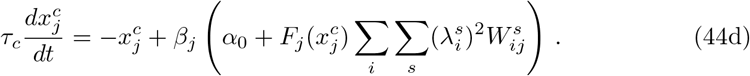

Because of Eq. (44c), at equilibrium all the sister mitral cells (all the 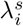) have the same value. Then, assuming, as above, that at *t* = 0 the average periglomerular activity is zero, it’s easy to see that the sister mitral cells have the same equilibrium values as they do in Eq. (42) if

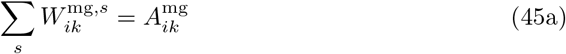

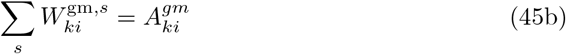

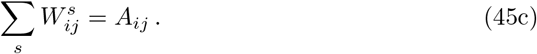

Thus, sister mitral cells can be used in more complicated models than the purely linear one we considered here.

## 3 Discussion

Most neural implementations of probabilistic inference require dense or all-to-all connectivity between elements, so that the explanatory power of all latent variables can be correctly accounted for. In common sensory settings where inference over hundreds of thousands of latent variables is not uncommon, such connectivity can require individual neurons to connect to hundreds of thousands of others, which is biologically implausible. In this work we have taken inspiration from the vertebrate olfactory system to show how such inference problems can be solved using sparse connectivity.

Naive olfactory inference would require each mitral cell to connect to hundreds of thousands of granule cells. However, in mice there are at least ~25 sister cells per glomerulus [22,23], and we showed that the sisters can share the connectivity, resulting in a substantial reduction in the required number of synapses per mitral cell. However, this sharing of connectivity comes at a cost: it requires coordination by a neural population. We have identified that population as periglomerular cells, based on their pattern of connectivity.

Our approach is not limited to the particular inference setting presented here. It can be applied to the more complex generative model in [2], and a large class of nonlinear models. As another example, our approach can be readily applied to the ‘sparse incomplete representations’ of [32], whose Eqs. (5) and (6) are directly analogous to our Eq. (10), and thus can be modified analogously to employ sparse connectivity.

### 3.1 Coordinating connectivity

One of the key requirements of our work is that sister cell connectivity matrices 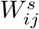 sum according to Eq. (14). This condition implies that the weights connecting a given granule cell *x_j_* to the sisters 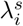 in glomerulus *i* sum to *A_ij_*. In the sparsest connectivity setting this requires that each granule cell connect to exactly one sister cell from each glomerulus. This may seem difficult to implement, but given the similarity in the temporal responses of sister cells, such sparse connectivity may be achievable through lateral inhibition among granule cell spines, as two spines each contacting sisters from the same glomerulus would receive very similar inputs, motivating the elimination of one to reduce redundancy. Nevertheless, although sparsifying connectivity was the motivation for our theory, the theory does not require it, as long as the condition in Eq. (14) is met. The main implication of this condition is that if a granule cell does contact multiple sisters from a given glomerulus, the weights are not redundant as eliminating one would disturb the weight condition and result in incorrect inference.

Perhaps a more fundamental issue is that we have assumed throughout that the correct connectivity – the affinity matrix *A_ij_* – is known. The key question is then how such ‘correct connectivity’ patterns can be learned, particularly in a setting where connectivity has to be coordinated across sister cells according to conditions like Eq. (14). Previous work [33] and current investigations [34] have examined this issue. However, neither of those studies used sister mitral cells to sparsify connectivity, and how such circuitry could be learned remains an open question.

### 3.2 Cortical readout

Although our work deals mainly with olfactory computation in the bulb, communicating the results of this computation to the cortex is obviously required. In Sec. 2.5 we outlined one scheme by which this could be done, in which cortical neurons effectively mirror the computation of olfactory bulb granule cells. If the projections from the mitral cells to the piriform cortex satisfy Eq. (40), the cortical readout at convergence would be an exact copy of the granule cell activations in the bulb. Needless to say, the main problem with such a scheme is how to satisfy this condition on the weights, and is likely made more difficult by the physical separation of the bulb and the cortex. One possibility is that the feedback connections from the cortex to the bulb could be used to learn the right connectivity. Another possibility is that an exact copy of the granule cell output is not required, and that random projections from the bulb to the cortex would suffice [35]. We leave the resolution of such issues to future work.

### 3.3 Predictions

Our model makes a number of experimentally testable predictions. Perhaps the easiest to test is the condition expressed in Eq. (36), which tells us that if periglomerular cells are perfect integrators, there is a nonlinear transformation of the mitral cell activity such that at convergence, the ratio of transformed sister mitral cell activity is independent of odour. This prediction breaks down if the periglomerular cells are leaky. However, at least in the linear case, the effect of leak is small (Figs. 9, top two panels, and Fig. 10C). It is admittedly hard to define ‘convergence’ in animals that are subject to periodic olfactory input due to the breathing cycle. Nevertheless, we can perhaps approximate it by the activity at the end of each breathing cycle in an anesthetized animal presented with an odour, and after several breathing cycles have elapsed. Our theory would then predict that, after a nonlinear transformation (which would have to be estimated from data), the ratio of the firing rates of two sister cells measured at this point would be the approximately the same regardless of odour, and would only depend on the identity of the sister cells.

A key computation required by our model is the (potentially weighted) average of sister mitral cell activity, temporally averaged and used to excite the sisters in a glomerulus. A variety of neural implementations of this computation are possible. Whatever its neural implementation, deactivating this component of the circuit should eliminate sister cell coordination and push the results of inference away from the MAP solution, and likely produce widespread, low amplitude activation of granule cells.

Although maximum sparsity is not required of our model, (see e.g. Fig. 3B) if, as we propose, the function of the sister mitral cells is to sparsify the mitral-granule cell connectivity required for inference, then maximally sparse connectivity solutions would be expected, in which granule cells contact at most one sister cell per glomerulus. Connectomics studies in the olfactory bulb should be able to tell us how often granule cells receive input from two or more sister mitral cells. Whether the connectivity is sparse or not, our theory requires that weights satisfy conditions like Eq. (14) for correct inference to occur. Hence, even if granule cells are found to contact multiple sisters, these connections are not redundant, and our theory would predict that experimentally perturbing or eliminating them would reduce inferential accuracy.

### 3.4 Summary

In summary, taking inspiration from the sister mitral cells in the vertebrate olfactory bulb, we showed how inference that typically requires dense connectivity can be performed using sparse connectivity. This means computations that would normally required hundreds of thousands of connections can be performed with a fraction of that. To the best of our knowledge our work is the first to propose a computational role for the sister mitral cells, and it makes a number of experimentally testable predictions. Despite its olfactory origins, our approach is quite general, and can be applied in other inference models and sensory modalities.

## 4 Methods

Here we provide additional details about the analyses used in the Main Text. In Sec. 4.1 we introduce a Lagrangian for the non-leaky (*ε* = 0) system, and modify it for the leaky case to gain insight into the resulting dynamics. In Sec. 4.2 we perform a linear analysis to determine how the various parameters influence the transient response properties of the system. Finally, in Sec. 4.3 we provide additional details about our simulations of the system dynamics.

### 4.1 Leaky periglomerular cells

To gain insight into the effect of pergilomerular leak, we start with a Lagrangian for the non-leaky setting

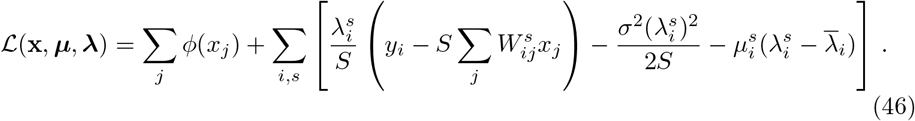

It is straightforward to show that

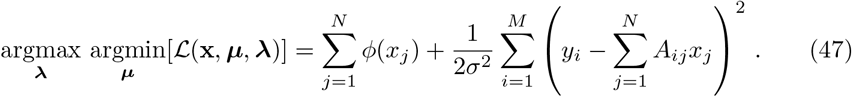

Consequently, if we minimize the Lagrangian with respect to **x** and ***μ*** and maximize it with respect to ***λ***, we recover the MAP solution. The resulting equations are

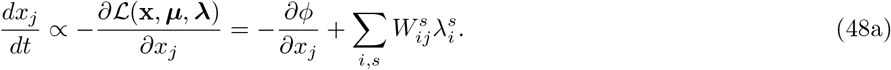

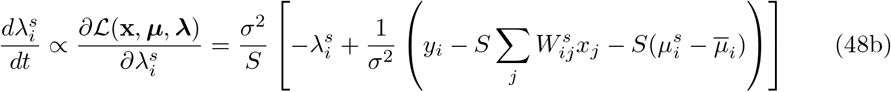

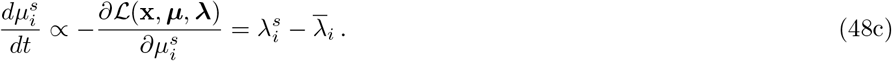

These equations correspond, up to scaling factors, to Eqs. (23), (20), and (19), respectively, except that the second equation above includes the additional term 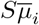. That term is, in fact, necessary to yield the correct dynamics. We dropped it because we can instead choose the initial conditions so that 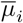 is zero, and under the periglomerular dynamics it will stay zero forever. Thus the circuit dynamics can be viewed as finding the saddle point of the Lagrangian of a constrained optimization problem derived from the original MAP objective.

To gain insight into the effect of the periglomerular leak we add a term proportional to 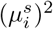 to the Lagrangian in Eq. (46),

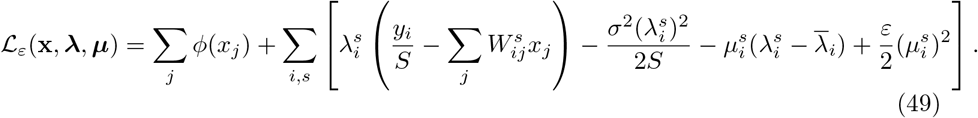

As is not hard to show, this introduces a term 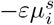 to the right hand side of Eq. (48c), which is the same as the leak term in Eq. (33b). To give us a Lagrangian that depends solely on **x**, we eliminate the auxiliary variables 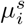 and 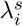. We start by minimizing 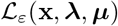 with respect to 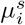, yielding

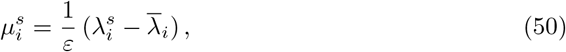

so that

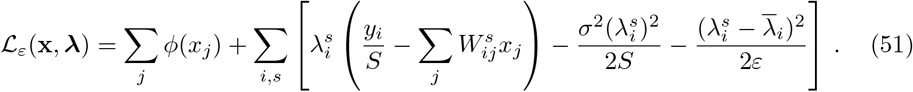

The final term couples the 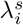 to their mean value computed over sisters. Extremizing with respect to 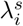 yields

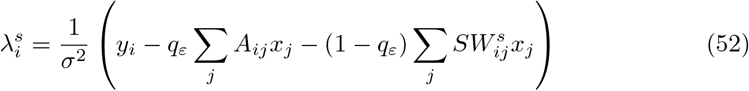

where

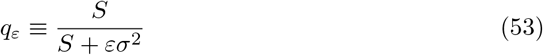

and we used, as usual, the constraint on the weights, Eq. (14). We can now insert this into Eq. (51) to derive a Lagrangian that depends only on **x**. Doing that is straightforward, although somewhat algebra-intensive, but it ultimately yields Eq. (26).

### 4.2 Linear dynamics

In addition to the number of sister cells *S*, our model has several parameters that affect the transient dynamics. To understand these effects, we performed a linear analysis of the model in Eq. (33). Our aim is a qualitative understanding, so we frequently opt for approximations that yield simple and tractable results over exact solutions.

We perform the linear analysis around the steady-state solutions to Eq. (33). Because of the threshold nonlinearity in Eq. (33d) only a small number of granule cell activations *x_i_* will be non-zero. We take the deviations around steady-state small enough so that the composition of this active set does not change. Adopting notation where vectors are in sister cell space, we write

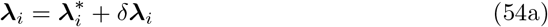

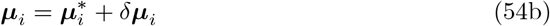

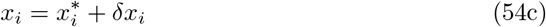

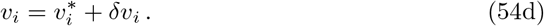

Here starred quantities are steady-state solutions to Eq. (33); quantities with a *δ* in front of them are infinitesimally small, and obey linear dynamics

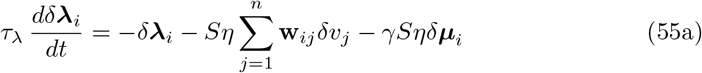

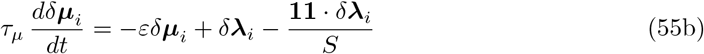

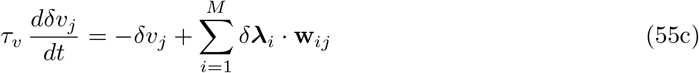

where

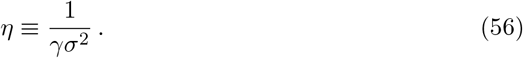

Because these are linear equations, they (generically) have solutions whose temporal part is given by *e^ξt^*. Consequently, derivatives with respect to time can be replaced by *ξ*, leading to

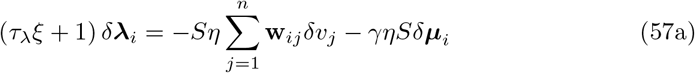

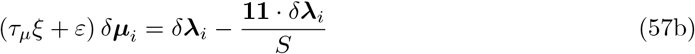

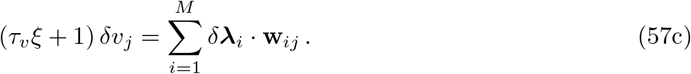

Our approach is to transform this set of equations to an eigenvalue equation in a single variable. To that end we eliminate *δ_**μ**_i__* and *δv_j_*, leaving us, after a small amount of algebra, with

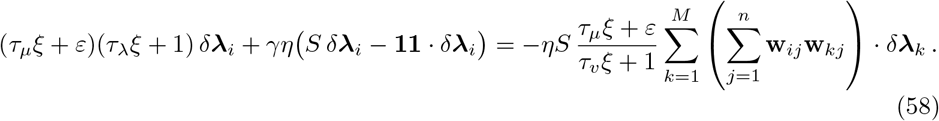

The *S* × *S* term in parentheses on the right hand side is the (*i, k*)^th^ block of an *MS* × *MS* matrix of rank *n*, whereas the set of vectors *δ**λ**_k_* contain *MS* components (*k* runs from 1 to *M* and *δ**λ**_k_* is an *S*-dimensional vector). Consequently, so long as *n* < *MS* (the regime we consider), that rank n matrix has two different classes of eigenvectors: those with zero eigenvalue and those with non-zero eigenvalue. We consider the former first.

For eigenvectors with zero eigenvalue, the left hand side of Eq. (58) must be zero. For that to happen there are, naively, two possibilities: *δ**λ**_i_* ∝ **1**, in which case *ξ* obeys

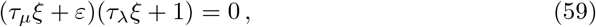

and *δ**λ**_i_* · **1** = 0, in which case *ξ* obeys

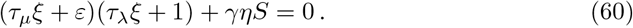

Again naively, this should result in four eigenvalues so long as *MS* > *n*.

To make this rigorous – and to uncover exactly when the above eigenvalues apply (as the naive conclusions are not quite correct) – we need a more involved analysis. Before proceeding with the general case, however, we note that there’s a special case: *τ_μ_ξ*+*ε* = 0, since in that case the right hand side of Eq. (58) is identically zero. Examining Eq. (57b) we see that when this holds, *δ**λ**_i_* ∝ **1**, which then determines *δv_j_* by Eq. (57c), and *δ**μ**_i_* by Eq. (57b). Thus there are M modes, corresponding to the root of Eq. (59); for these,

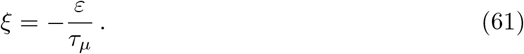

When *τ_μ_ξ* + *ε* ≠ 0, there is an (*MS* – *n*)-dimensional space of vectors, denoted 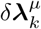, that obey

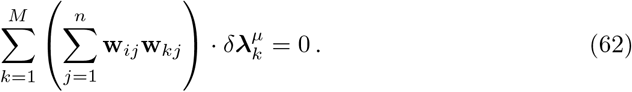

We need to choose a linear combination of these for which the left hand side of Eq. (58) is zero; that is, we need to choose a set of *a_μ_* such that (after rearranging terms slightly)

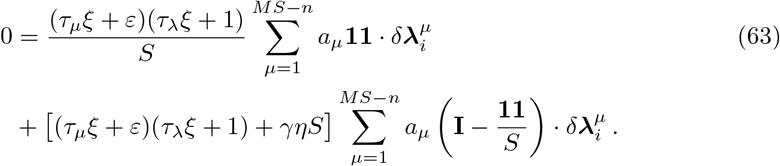

Since this equation must be satisfied for all *i*, there are MS equations (*i* runs form 1 to *M* and 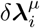 is an *S*-dimensional vector). But there are only *MS* – *n* adjustable parameters, so in general the only solution has all the *a_μ_* = 0. There are, though, two ways to find nontrivial solutions. One is to set the first term in parentheses to zero (i.e., enforce Eq. (59)). Then, because 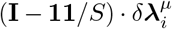 spans *S* – 1 dimensions, Eq. (63) corresponds to *M*(*S* – 1) equations. Because there are *MS* – *n* adjustable parameters, there is a nontrivial solution if *MS* – *n* – *M*(*S* – 1) > 0; that is, if *M* > *n*. Note that because we have already taken into account the solution with *τ_μ_ξ* + *ε* = 0, these solutions must have *τ_λ_ξ* + 1 = 0. The other way to find nontrivial solutions is to set the term in brackets in Eq. (63) to zero (i.e., enforce Eq. (60)). Then, because 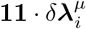 spans one dimension, Eq. (63) corresponds to *M* equations, and so there is a nontrivial solution if *MS* – *n* – *M* > 0; that is, *M*(*S* – 1) > *n*.

In summary, when the right hand side of Eq. (58) is zero, we have the following eigenvalues, all of which are exact: *M* modes with

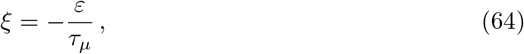

[*M* – *n*] ^+^ modes with

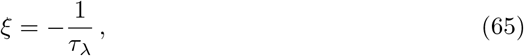

and [*M* (*S* – 1) – *n*]^+^ modes with

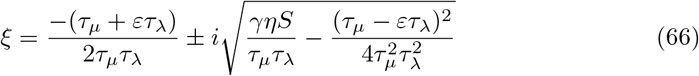

where 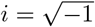 (when it’s is not an index). Because *ξ* can take on two values, there are 2[*M*(*S* – 1) – *n*]^+^ modes of this type.

Two comments are in order. First, when there is only one sister cell (*S* = 1), the mode in Eq. (66) does not exist, as that mode requires *M*(*S* – 1) > *n*. Second, for the modes given in Eq. (64), (65) and (66), 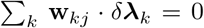; this in turn implies, via Eq. (57c), that *δv_j_* = 0. Thus, these modes involve the periglomerular and mitral cells, but not the granule cells.

When the right hand side of Eq. (58) is nonzero, analysis in *δ**λ**_i_* space is difficult. However, we can instead work in *δv_j_* space: eliminating *δ**λ**_i_* and *δ**μ**_i_* from Eq. (57), we can write down an eigenvalue equation for *δv_j_*; after tedious but straightforward algebra, including application of the Sherman-Morrison formula to invert the operator on the left-hand side of Eq. (58), we arrive at

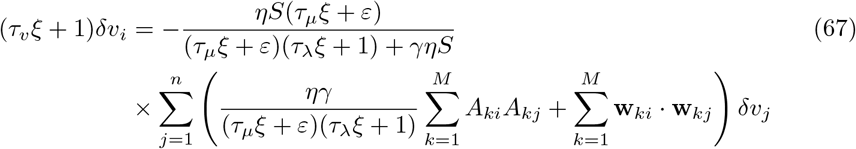

where we used Eq. (14) to write **1** · **w**_*kj*_ = *A_kj_*.

Finding exact non-trivial solutions to Eq. (67) requires finding the eigenvalues of the sum of the two matrices on the right hand side of this equation. That’s difficult in general, so instead we make an approximation: we replace the second matrix with the identity. We justify this by arguing that its eigenvalues are narrowly distributed around 1.

To show that, we start by writing

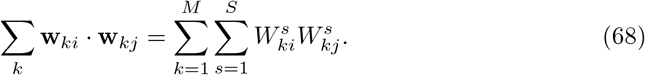

For a given *k* and *j*, The elements of 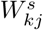 are nonzero for only one value of *s* and zero for the rest (see Eq. (18)). Consequently, they are not *iid*, which makes it difficult to compute the eigenvalue spectrum. However, a reasonable approximation to 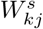 is

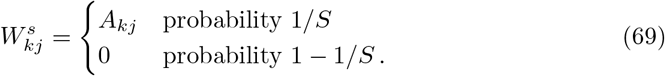

In that case, 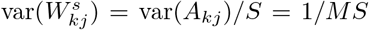, implying that 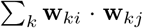 approximately follows a Marcenko-Pastur distribution with parameters (1, *n/MS*) [36]. For this distribution, the eigenvalues lie in the range

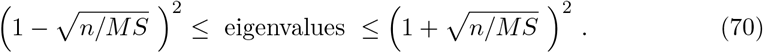

When *n* ≪ *MS*, the regime of interest, these eigenvalues are very narrowly distributed around 1. Thus, the matrix in Eq. (68) is, to good approximation, the identity. This approximation breaks down as *n* increases, but we’re mainly interested in small *n*, so that is not a problem.

With this approximation, the only nontrivial matrix left in Eq. (67) is the one involving the *A_kj_*. The elements of *A_kj_* are drawn *iid* from 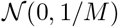, so

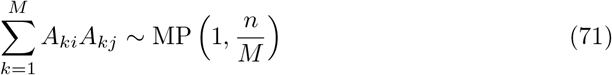

where MP denotes the Marchenko-Pastur distribution. Using *ν* to denote an eigenvalue of this distribution, we see that Eq. (67) can be approximated as

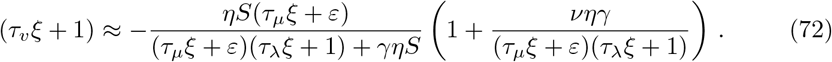

There are *n* eigenvalues, corresponding to the fact that *j* runs from 1 to *n* in Eq. (67), so there are *n* sets of solutions to this equation. (We say “sets of solutions”, rather than just one, because Eq. (72) is a polynomial in *ξ*, which has several roots.) The positive eigenvalues lie in the range

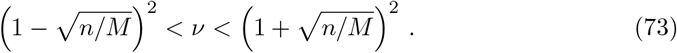

If *n* < *M*, all of the eigenvalues lie in this range, while if *n* ≥ *M*, only *M* eigenvalues lie in this range; the other *n* – *M* are zero.

#### 4.2.1 Approximate solutions

To solve to Eq. (72), our first step is to write it is a polynomial,

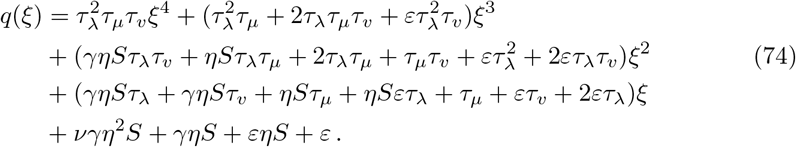

We’re looking for values of *ξ* that satisfy *q*(*ξ*) = 0. Note that *q*(*ξ*) depends on *ν*, which means solutions to *q*(*ξ*) = 0 will also depend on *ν*; we drop that dependence to reduce clutter.

Because *q*(*ξ*) is quartic, an exact analytic expression for its roots is available, but it is too complex to yield insight. Instead, we take a perturbative approach, which rests on the observation that *η* is large, on the order of 100 (see its definition, Eq. (56) and Table 1). To take advantage of this, we scale *ξ* by a factor of 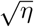. Choosing a scaling that gives us a dimensionless quantity, we make the change of variables

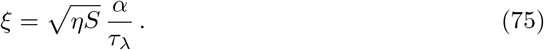

Then, defining the time constant ratios

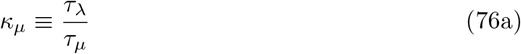

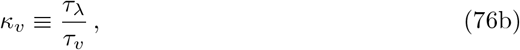

and working to first order in 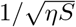 we find, after straightforward algebra, that *q*(*ξ*), expressed in terms of *α* (and denoted, in a slight abuse of notation, *q*(*α*)) is given approximately by

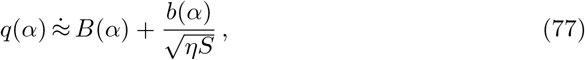

where 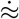 indicates approximate equality up to multiplicative constant and

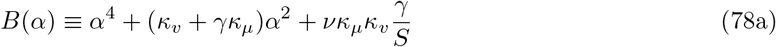

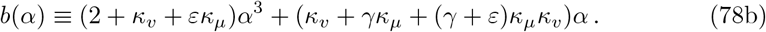

In the large *η* limit, the roots of *q*(*α*) are determined by those of *B*(*α*). Defining

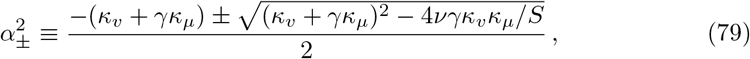

the corresponding four roots are ±*iα*_±_. The argument of the square root is (*κ_v_* – *γκ_μ_*)^2^ + 4*γκ_v_κ_μ_*(1 – *ν/S*), which is guaranteed to be positive if *ν* < *S*. From Eq. (73), this requires 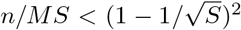. We’ll restrict ourselves to this regime, which ensures that both 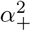 and 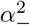 are negative, which in turn means all four of these roots are purely imaginary.

To compute the corrections to these roots, we let *α* = *α*_0_ + *α*_1_ where *α*_0_ is any of the above four roots. Then, performing a Taylor expansion of *q*(*α*), Eq. (77), around *α*_0_, we have

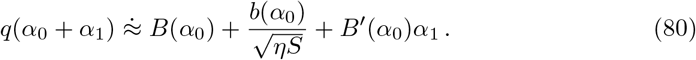

Setting this to zero and solving for *α*_1_ gives

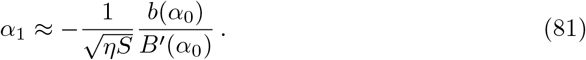

Using Eq. (78) for *B*(*α*_0_) and *b*(*α*_0_), setting 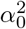 to 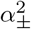, and using Eq. (79) to simplify the resulting expression, we arrive at

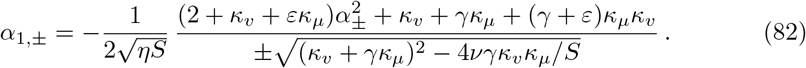

We thus have (using Eq. (75))

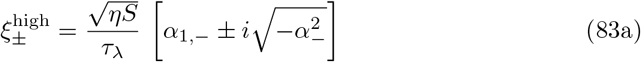

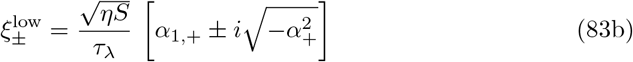

with *α*_1,±_ given in Eq. (75) and 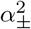 given in Eq. (79). The “high” and “low” superscripts refer to the fact that 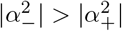, as is easy to see from Eq. (79).

It is instructive to consider the large *S* limit, which greatly simplifies the roots. Focusing first on the high frequency roots, in this limit we have

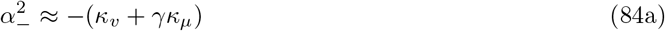

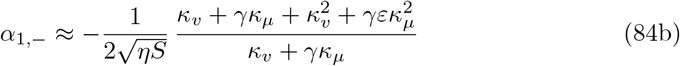

so that, using Eq. (75),

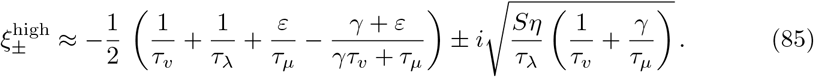

To approximate the low frequency roots, which correspond to 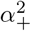, we perform a Taylor expansion of the square root in Eq. (79) around (*κ_v_* + *γκ_μ_*)^2^, yielding

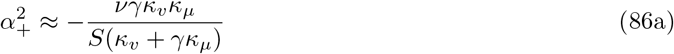

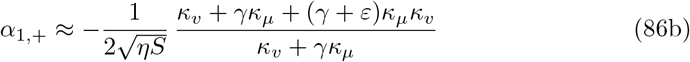

so that, using Eq. (75),

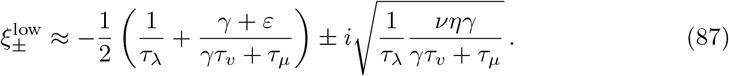

In summary, our goal was to find solutions to Eq. (74) for each value of *ν*, where *ν* is drawn from the Marchenko-Pastur distribution MP(1, *n/M*). This implies that there is a distribution of solutions, *ξ*, which we can find by solving for *ξ* at each *ν*. We did that perturbatively, yielding the high and low frequency solutions given in Eq. (83) (with approximate expression for these quantities given in Eqs. (85) and (87)). Note that if *ν* = 0 (which can happen when *n* > *M*), *α*_+_ = 0 (see Eq. (79)). When that happens, 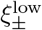 takes on only one value, not two (see Eq. (83b)), and so there are three possible solutions.

The number of modes when the right hand side of Eq. (58) is nonzero, then, depends on *n*. There are always 2n high frequency modes. When *n* ≤ *M* (so that *ν* is strictly positive), there are also 2*n* low-frequency oscillatory modes. When *n* > *M*, on the other hand, *n* – *M* of the eigenvalues, *ν*, are zero, and the rest are positive, resulting in *n* – *M* decaying modes and 2*M* low-frequency oscillatory modes. We thus have 2*n* – [*n* – *M*] ^+^ decaying and low-frequency oscillatory modes, for a total of

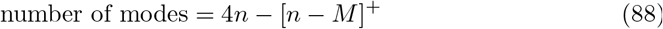

for Eq. (67).

All modes of the system are tabulated in Table 2. They are given exactly by Eqs.(64), (65) and (66), and approximately by (83). All of these modes have a decay associated with them, and the latter three also have oscillation frequencies. For simplicity, we considered the large S limit, so we used Eqs. (85) and (87) for the approximate modes given in Eq. (83). Assuming *M*(*S* – 1) > *n*, the total number of modes is *M* + [*M* – *n*]^+^ +2(*M*(*S* – 1) – *n*) + 4*n* – [*n* – *M*]^+^. Adding these together gives 2*MS* + *n*, as it should.

**Table 2.** Linear analysis modes and their properties. The last two modes correspond to the large *S* limit of Eq. (83).

| $\vert \text{Re}(\xi)\vert ^{-1}$: Decay time constant | $\vert \text{Im}(\xi)\vert $: Oscillation frequency | Number of modes | Source |
| --- | --- | --- | --- |
| $\frac{\tau_\lambda}{\varepsilon\kappa_\mu}$ | 0 | $M$ | Eq. (64) |
| $\tau_\lambda$ | 0 | $[M - n]^+$ | Eq. (65) |
| $\frac{2\tau_\lambda}{1 + \varepsilon\kappa_\mu}$ | $\frac{1}{\tau_\lambda} \sqrt{\gamma\eta\kappa_\mu S}$ | $2[M(S - 1) - n]^+$ | Eq. (66) |
| $\frac{2(\gamma\kappa_\mu + \kappa_v)\tau_\lambda}{\kappa_v^2 + \gamma\varepsilon\kappa_\mu^2 + \gamma\kappa_\mu + \kappa_v}$ | $\frac{1}{\tau_\lambda} \sqrt{\eta(\gamma\kappa_\mu + \kappa_v)S}$ | $2n$ | Eq. (85) |
| $\frac{2(\gamma\kappa_\mu + \kappa_v)\tau_\lambda}{(\gamma + \varepsilon)\kappa_\mu\kappa_v + \gamma\kappa_\mu + \kappa_v}$ | $\frac{1}{\tau_\lambda} \sqrt{\frac{\nu\eta\gamma\kappa_v\kappa_\nu}{\gamma\kappa_\mu + \kappa_v}}$ | $2n - [n - M]^+$ | Eq. (87) |

#### 4.2.2 Stability

The perturbative corrections in Eq. (82) allow us to assess the stability of the linearized dynamics. For stability, both *α*_1,+_ and *α*_1,−_ (which are real) must be negative. Combining Eq. (79) with Eq. (82), we see that this gives us the two conditions,

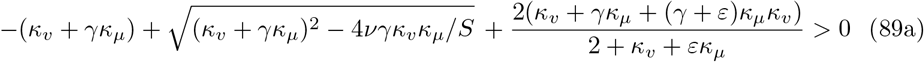

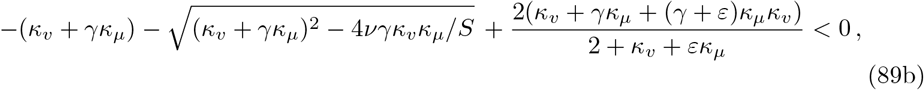

which can be simplified to just one,

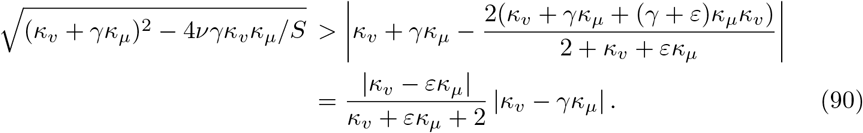

As above (see comments following Eq. (79)), in the regime of interest, 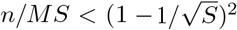, the left hand side of this equation is greater than |*κ_v_* – *γκ_μ_*|. The ratio on the right hand side is less then 1, so the right hand side is less than |*κ_v_* – *γκ_μ_*|. Consequently, this inequality is satisfied. Thus, at least in the large *η* limit, all roots are stable.

### 4.3 Simulations

The base values of all parameters used in simulations are listed in Table 1. To model each granule cell connecting to a single sister cell from each glomerulus, we selected for each glomerulus *i* and granule cell *j* a random sister cell; see Eq. (18). The non-zero concentrations of the presented odours were set to 1, except in Figures 5, 6, 9 and 12, where the true odour had the default number *n* = 3 non-zero components but at concentrations of 0.8, 1, and 1.2, to aid visual assessment of convergence to the MAP solution.

To assess the variability of the various response characteristics we usually chose to present the same odour to different random instances of the olfactory bulb, rather than picking different odours within the same olfactory bulb. Thus all references to trials are to the same odour presented to different olfactory bulbs unless stated otherwise.

The quantities in Fig. 8 were computed as follows. Amplitude spectra for panel A and panel B were computed as the absolute value of the Fourier transform of the mitral cell responses in the time interval *t* = 0.3 — 0.6 seconds, averaged over all mitral cells and 20 trials. The decay time constants in panel C were computed from the slope of a linear fit to time of the log RMS error of the mitral cell activations relative to their final value, for the interval *t* = 0.4 – 0.6 seconds following odour onset averaged over 20 trials.

For all simulations we used the forward Euler method with a time step of 10^−3^ ms. To confirm that our networks performed MAP inference, we compared solutions to those found by the convex optimization package CVXPY [37] using the splitting conic solver (SCS), with `eps` set to 5 × 10^−13^, applied to the MAP problem in Eq. (7) expressed as the constrained optimization

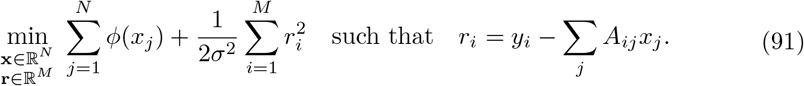

The code used to run the simulations and produce the figures are available at https://github.com/stootoon/sister-mcs-release.

## Acknowledgments

We thank members of the Gatsby Unit and the Latham and Schaefer labs for useful discussions. This work was supported by the Gatsby Charitable Foundation, Wellcome Trust Investigator grants 110174/Z/15/Z (A.T.S.), 110114/Z/15/Z (P.E.L.), and by the Francis Crick Institute, which receives its core funding from Cancer Research UK (FC001153), the UK Medical Research Council (FC001153), and the Wellcome Trust (FC001153). For the purpose of Open Access, the authors have applied a CC BY public copyright licence to any Author Accepted Manuscript version arising from this submission.

